# Involvement of the cellular prion protein in seeding and spreading of sarkosyl-derived fractions of Alzheimer’s disease in *Prnp* mutant mice and in the P301S transgenic tauopathy mice model

**DOI:** 10.1101/2024.01.20.576414

**Authors:** Julia Sala-Jarque, Vanessa Gil, Pol Andrés-Benito, Laia Lidón, Renato Eduardo Yanac-Huertas, Clara F López-León, Félix Hernández, Jesús Ávila, José Luis Lanciego, Jordi Soriano, Mario Nuvolone, Adriano Aguzzi, Rosalina Gavín, Isidro Ferrer, José Antonio del Río

## Abstract

The natural cellular prion protein is known to play several roles during development and adult brain. Far from its pathological roles in prionopathies, the non-pathogenic cellular prion protein has been described as a receptor for several amyloid in oligomeric and prefibrillar forms. For some amyloids, specific domains of the protein play a crucial role in modulating amyloid’s cellular uptake and seeding properties. In most studies, the functions and the role of putative amyloid receptors have been analyzed by using brain extracts derived from human neurodegenerative patients. Another strategy has been to modify the genetic dosage of the natural prion protein in genetic models of different diseases. In this study, we take advantage of both approaches to examine whether this protein plays a role in the seeding and spreading of pathogenic tau. Our results point to a role of the natural prion protein in the emergence of pathogenic tau in a mouse model overexpressing the mutation P301S of the human tau gene. In contrast, its role is minor when sarkosyl-derived brain samples of Alzheimer’s disease are used. In fact, our results indicate that the use of this type of sample is not adequate to determine the role of a putative receptor in tau seeding and spreading.

## Introduction

Human tauopathies are clinically, biochemically, and heterogeneously neurodegenerative diseases characterized by the deposition of the abnormally misfolded and aggregated microtubule-associated protein tau in specific brain regions (i.e., [20, 64, 72, 118]). Each tauopathy has an exclusive clinicopathologic phenotype defined by: i) the types of tau deposits from exon 10 splicing (3Rtau or 4Rtau); ii) the aggregation status of ptau species (puncta/granular/diffuse/fibrillar/tangles/pretangles); and iii) the affected brain regions and cells, the latter including neurons, astrocytes and oligodendrocytes (e.g., [46, 64, 72, 108, 110, 118]). Tau aggregates are found in sporadic and familial Alzheimer’s disease (AD), certain prion diseases as in Gertsmann-Straüsler-Scheinker (GSS); and in pure forms, either sporadic as in primary age-related tauopathy (PART), aging-related tau astrogliopathy (ARTAG), progressive supranuclear palsy (PSP), argyrophilic grain disease (AGD), and glial globular tauopathy (GGT), or familial, linked to mutations in *MAPT* gene: familial frontotemporal lobar degeneration-tau (FTLD-tau) (e.g., [27, 50, 54, 64, 72-74, 89, 96, 101]. Most human tauopathies are characterized by exhibiting a transregionally spreading proteinopathy resulting in stereotyped spatiotemporal progression patterns [10, 40, 59]. Thus, after the inoculation of ptau-containing samples in mice brains, pathologic ptau aggregates emerged both at the injection sites and in connected brain regions (e.g., [34, 41]). Thus, this approach has been used to elucidate the mechanisms of cell uptake, seeding and brain propagation of different tau species (oligomeric or fibrillar) as well the role of the endogenous tau in these processes (e.g., [8, 48, 51, 52, 90]). However, a complete explanation of these processes is unavailable for all tau isoforms and different tauopathies [64] especially when affected cells are other than neurons [45]. From the different mechanisms of uptake of misfolded proteins (see [41]), receptor-mediated uptake is well established. In this respect, a recent study suggests the presence of shared putative neuronal receptors for tau, Aβ, and α-synuclein [98]. In fact, among others, the low-density lipoprotein receptor-related protein 1 (LRP1) [1, 104], the cellular prion protein (PrP^C^) [26, 35, 79] or the lymphocyte-activation gene 3 (LAG3) [30] have been revealed as functional neuronal receptors of tau oligomers in some experimental models. PrP^C^ is a cell surface GPI-anchored protein expressed in several tissues with especially high levels expressed by neurons and glial cells [2, 21, 83, 93]. PrP^C^ is known for its crucial role in the pathogenesis of human and animal prionopathies [3, 13, 102]. Recent increasing knowledge about the participation of PrP^C^ in prion pathogenesis contrasts with the puzzling data regarding its natural physiological role probably related to its specific interactions [57, 60, 78, 84, 129, 131]. The PrP^C^ central region contains a central hydrophobic domain (HD or HR, aa 110/113-133) and a charged cluster domain (CD, aa 94–110), both involved in binding with different oligomeric/fibrillar amyloids (i.e., scrapie prions and Aβ or α-synuclein, respectively) [44, 105, 106] reviewed in [41]). From histopathological studies, we know that PrP^C^ is localized in dystrophic neurites and amyloid plaques in advanced AD [122, 123, 137]. Thus, although the physiological role modulating neuronal activity of amyloid-PrP^C^ interaction has been described (see for Aβ [77], α-synuclein [44, 125] or tau [98]), their participation in tau seeding and spreading is still elusive and very few data is based on the role of PrP^C^ in a genetic model of AD [119]. Indeed, by using a double mutant mouse of APP and *Mapt* the authors point out that the absence of PrP^C^ shows a moderate decrease in AT8 pTau deposits, but no changes in Aβ accumulation [119]. In fact, three recent studies that overexpressed PrP^C^ might bind to tau but their putative role in tau seeding and spreading are not analyzed [35, 39, 79]. However, due to an additional study pointing out that PrP^C^ cannot bind to α-synuclein oligomers [75], which contrasts with other studies [12, 44, 109, 125, 128], we consider that these putative PrP^C^-tau interactions *in vivo* merit further research. Thus, in this study, we aimed to explore the role of PrP^C^ in seeding and spreading *in vivo* of pathogenic tau derived from sarkosyl-insoluble samples of AD and the putative blockage of spreading of the pathogenic tau by using a soluble form of the full-length PrP^C^ to block cellular tau uptake and transmission. To develop these aims, we took profit of mutant *Prnp* mice overexpressing the GPI truncated form of tau: the “anchorless” GPI^−^ *Prnp* (Tg44) mice strain [31]. In addition, and to compare the observed results, the effect of the genomic deletion of the *Prnp* gene in P301S mutant mice is analyzed and the putative changes in ptau generation after inoculation of adeno-associated virus (AAV) expressing human mutated tau at P301L in *Prnp^0/0^ vs* wild type is also analyzed. Obtained results point to a minor role of PrP^C^ in the seeding and spreading of sarkosyl-insoluble samples of AD but a role after genetic ablation in P301S mice. Lastly, we demonstrate that the overexpression of the truncated form of PrP^C^ is unable to impair seeding of ptau in sarkosyl-insoluble inoculated mice.

## Materials and Methods

### Mice strains

Adult ZH3 *Prnp^0/0^* mice line was generated by A. Aguzzi (Switzerland) (see [97] for details). In addition, mice overexpressing the secreted form of PrP^C^ lacking their GPI anchor (GPI^−^ *Prnp* (Tg44) mice, kindly provided by Vincent Beringue, INRA UR892, Virologie Immunologie Moléculaires, Paris, France) were used [31]. Moreover, mice lacking *MAPT* [37](kindly provided by Jesús Ávila, CBM-UAM, Madrid) were also used. In addition, P301S (PS19) transgenic mice [5] (Jackson Laboratories ref: 008169) that overexpress the human mutation of tau under the cellular prion protein promoter [133], some Nestin-cre/*Prnp^flox/flox^* mice generated in our laboratory by crossing *Prnp*^flox/flox^ [126] and Nestin-cre [43] were also used. A total of 74 adult (2-5 months old) male mice (*Prnp^+/+^*= 30, *Prnp^0/0^* = 12, GPI^−^ *Prnp* = 11, *Mapt^0/0^* = 5, P301S =11 and Nestin-cre/*Prnp^flox/flox^*mice = 5 were used in the present study. All the animals were kept in the animal facility of the Faculty of Pharmacy at the University of Barcelona under controlled environmental conditions and were provided food and drink *ad libitum*. All experiments were performed under the guidelines and protocols of the Ethical Committee for Animal Experimentation (CEEA) of the University of Barcelona, and the protocol for the use of animals in this study was reviewed and approved by the CEEA of the University of Barcelona (CEEA approval #276/16 and #141/15).

### Human brain samples

A total of 10 sarkosyl-insoluble fractions from the frontal cortex (area 8) of AD patients were obtained and characterized. After biochemical and histological analysis, we selected one AD sample from a female patient of 56 years (Braak stage VI), since no additional lesions (ischemia or other amyloid aggregation) were detected after the conventional neuropathological analysis. Brain samples of the AD patients were obtained from the Institute of Neuropathology Brain Bank, Bellvitge University Hospital, following the guidelines of the Spanish legislation on this matter (Real Decreto Biobancos 1716/2011) and the approval of the local ethics committee of Bellvitge University Hospital (Hospitalet de Llobregat, Barcelona, Spain). The agonal state was short with no evidence of acidosis or prolonged hypoxia; the pH of each brain was between 6.8 and 7. At the time of autopsy, one hemisphere was fixed in paraformaldehyde and the other hemisphere was cut into coronal sections 1 mm thick, and selected brain regions were dissected, immediately frozen at −80°C, put on labeled plastic bags, and stored at −80°C until use; the rest of the coronal sections were frozen and stored at −80°C following the protocols published in [7]

### Sarkosyl-insoluble fraction preparation

Brain tissue was weighed and then homogenized using a Dounce homogenizer in 10 volumes of fresh homogenization buffer [0.8 M NaCl, 1 mM EGTA, 10% sucrose, 0.01 M Na_2_H_2_P_2_O_7_, 0.1 M NaF, 2 mM Na_3_VO_4_, 0.025 M β-glycerolphosphate, 0.01 M Tris-HCl pH 7.4] containing protease inhibitors (Roche, Switzerland). For sarkosyl preparations, after centrifugation at 16,000 rpm for 22 min at 4°C, the supernatant was reserved (SN1). The pellet was resuspended in 5 volumes of homogenization buffer and centrifuged again at 14,000 rpm for 22 min at 4°C. The resulting supernatant (SN2) was then combined with the SN1, and the mixture (SN1 + SN2) was incubated with 0.1% N-lauroyl sarcosinate (sarkosyl; Sigma-Aldrich) and placed on a rotating shaker for 1 h at room temperature. The mixture was centrifuged at 35,000 rpm for 63 min at 4°C. Next, the supernatant was discarded, and the remaining pellet (sarkosyl-insoluble fraction) was washed and resuspended in 50 mM Tris-HCl, pH 7.4 (200 µl/g starting material). Finally, 100 µl aliquots were stored at −80°C until use. Protein concentrations were determined using the Pierce™ BCA assay kit (Sigma-Aldrich), and equal amounts of protein were analyzed by western blot.

### Biochemical analysis

Samples were characterized by dodecyl sulfate-polyacrylamide gel electrophoresis (SDS-PAGE), followed by western blot. Protein extracts were boiled at 100°C for 10 min, followed by 6% SDS electrophoresis, and were then electrotransferred to nitrocellulose membranes for 1 hour at 4°C. Membranes were then blocked with 5% fat milk in 0.1M Tris-buffered saline (pH 7.4) for 1 hour and incubated overnight in a 0.5% blocking solution containing primary antibodies. After incubation with peroxidase-tagged secondary antibodies (1:2000 diluted, Sigma-Aldrich), membranes were revealed with an ECL-plus chemiluminescence western blot kit (Amershan-Pharmacia Biotech).

### Tau RD P301S Biosensor cell line experiments

The Tau RD P301S Tau biosensor [ATCC^®^ CRL-3275^TM^] was purchased from ATCC. Cells were grown in maintenance medium made of Dulbecco’s Modified Eagle Medium (DMEM; ThermoFischer Scientific) supplemented with 10% fetal bovine serum (FBS; ThermoFischer Scientific), 1% GlutaMax (Gibco), and 1% Penicillin/Streptomycin (ThermoFischer Scientific) in 75 cm^2^ culture flasks (Nunc). Cells were maintained at 37°C and 5% CO_2_ in a humidified incubator and passaged every 3 days when confluent. Tau biosensor cells were plated in 96-well poly-D-lysine (Sigma-Aldrich) (0.1 mg/ml) coated plates at a density of 35,000 cells/well (total volume 130 µl) in maintenance medium and cultured at 37°C in a 5% CO_2_ incubator overnight. The transduction mixture was prepared following the manufacturer’s protocol [65]. Briefly, 1.5 µl of the sample was combined with 8.5 µl of Opti-MEM medium (ThermoFischer Scientific). Next, a mixture of 1.25 µl of Lipofectamine-2000™ reagent (ThermoFischer Scientific) and 8.75 µl of Opti-MEM was then added to the sample mixture to a final volume of 20 µl and incubated for 1 h at room temperature. Mixtures with empty liposomes were included as negative controls. Tau biosensor maintenance medium was gently removed and replaced with 130 μl of pre-warmed Opti-MEM before 20 μl of the transduction mixture was added to the cells. Twenty-four hours later, cells were washed once with pre-warmed 0.1 M PBS and fixed with 4% phosphate-buffered paraformaldehyde (PFA) for 15 min at room temperature. Next, PFA was removed, and cells were washed three times 5 min each with 300 μl of 0.1 M PBS. Finally, 300 μl of 0.1 M PBS with 0.02% sodium azide was placed in each well, and the plate was sealed with Parafilm^TM^ and kept at 4°C until analysis. Using this reporter cell line the following samples were analyzed: Fibrillated or monomeric tau K18 at a final concentration of 0.01 µM (Bio-techno (ref: SP-496-100)). Human and murine α-synuclein pre-formed fibrils, both 0.1 µg/µl (a gift from Masato Hasegawa, Tokyo Metropolitan Institute of Medical Science, Japan). Monomeric tau Cy5 at a final concentration of 100 nM (kindly provided by Jesús Ávila, CBM-UAM, Madrid). Sarkosyl-insoluble fractions of P301S^+/−^ and P301S^−/−^ mice were used at a final concentration of 0.003 µg/µl. Sarkosyl-insoluble fractions from human brains were used at a final concentration of 0.003 µg/µl. In addition, full-length tau and phosphorylated full-length tau (human sequence, provided by Jesús Ávila, CBM-UAM, Madrid) were used at 100 and 200 nM in this study.

### Isolation of tau seeds generated in the Biosensor cell line

Tau biosensor cells were plated in 60-mm-diameter poly-D-lysine (0.1 mg/mL)-coated Petri dishes at a density of 200,000 cells/well in maintenance medium at 37°C and 5% CO_2_ in a humidified incubator. On the following day, cells were transduced using the same transduction protocol described in the previous section, always using sarkosyl-samples from P301S^+/−^ or P301S^−/−^ mice, and empty liposomes as the vehicle control condition. Twenty-four hours later, the transduction reaction was removed, and cells were washed once in prewarmed sterile 0.1 M PBS and further maintained with fresh maintenance medium. Forty-eight hours later, cells were washed twice with pre-warmed sterile 0.1 M PBS. Next, cells were scraped from the Petri dish surface in Triton X-100 lysis buffer (0.05% Triton X-100 in sterile 0.1 M PBS supplemented with 1X protease inhibitors) and pelleted at 500 x *g* for 5min at 4°C. Supernatants were then centrifuged at 1,000 x *g* for 5min at 4°C. Next, 10% of the supernatant volume of each sample was kept as the “Triton X-100 soluble fraction” (TSF). Importantly, each fraction was named based on the sample used in the transduction step of the Tau biosensor cells. Thereby, the brain homogenate from P301S^+/−^ mice produced “TSF-P+” fractions, the brain homogenate from P301S^−/−^ mice produced “TSF-P-“, and the empty liposomes (vehicle) condition produced “TSF-V.” The remaining TSF was centrifuged at 50,000 rpm for 30 min at 4°C. After that, the pellet was washed in 0.1 M PBS and further centrifuged at 50,000 rpm for 30 min at 4°C. The resulting pellet was resuspended in 50 µL of 50 mM Tris-HCl, pH 7.4 and reserved as “Triton X-100 insoluble fraction” (TIF), referred to as the “TIF-P+,” “TIF-P-,” and “TIF-V.” Aliquots of all fractions were stored at −80°C until further use. Protein concentrations were determined using the Pierce™ BCA assay kit (Sigma-Aldrich, Germany).

### Negative electron microscopy staining

For transmission electron microscopy (TEM) experiments, tau samples were fixed to carbon-forward-coated copper grids, and negative stained with buffered 1% uranyl acetate (pH 7.4). The samples were then placed in silica-based desiccant for a minimum of 2 h and examined with a Jeol JEM-1010 transmission electron microscope.

### Mouse tissue homogenation and preparation

The mouse brain tissue was weighted, and 1 ml of 0.1 M PBS supplemented with protease inhibitors (Roche, Switzerland) was used per 20 mg of tissue. The sample was homogenized for 10 min, using a Polytron^TM^, and 50 µl aliquots were then stored at −80°C until use. Protein concentrations were determined using a BCA assay, and equal amounts of protein were analyzed by western blot.

### Stereotaxic surgery

For surgery, mice were deeply anesthetized with isoflurane and placed in a stereotaxic apparatus (Kopf Instruments, USA). Unilateral stereotaxic injections were performed into the right hippocampus (AP: 1.4 mm from Bregma; LM: 1.5 mm; DV: 1.5 mm). 2.5 µl of sarkosyl extract dissolved in 100 mM Tris-HCl was inoculated, using a Hamilton syringe, into the upper corpus callosum/cortex at DV-1 mm. A similar protocol was used for the inoculation of AAV2r.*Syn*.P301L (VectorBuilder, Neu-Isenburg, Germany). Following the injection, the needle was kept in place for an additional 3 min before withdrawal. The surgical area was cleaned with sterile saline, and the incision was sutured. Mice were monitored until recovery from anesthesia and were checked regularly following surgery.

### Primary antibodies used in the study

The following primary antibodies were used in the study: monoclonal mouse anti-AT8 (1:50 dilution) (ThermoFischer Scientific, catalog no. MN1020), polyclonal rabbit anti-Tau-phosphoSer422 (pSer422) (1:75 dilution) (Life Technologies, catalog no. 44-764G), monoclonal mouse anti-6H4 (1:1000 dilution) (ThermoFischer Scientific, catalog no. 01-010), human tau-specific anti-Tau13 (1:200 dilution) (Biolegend, catalog no. 835201), murine tau-specific antibody anti-T49 (1:200 dilution) (Merck Millipore, catalog no. MABN827), mouse anti-human tau-specific HT7 (1:600 dilution) (ThermoFischer Scientific, catalog no. MN1000), monoclonal mouse MC-1 anti-tau (a gift of Prof. Peter Davis, [70]). The different tau isoforms 3R and 4R were checked with monoclonal RD3 (1:200 dilution) (clone 8E6/C11) and RD4 (1:200 dilution) (clone 1E1/A6) antibodies (Merck Millipore). Other antibodies used were rabbit anti P62/ SQSTM1 (1:100 dilution) (Progen, ref GP62-C), rabbit anti-GFP (1:500 dilution) (Invitrogen, catalog no. A11122), monoclonal mouse anti-Actin (1:1000 dilution) (Millipore, catalog no: MAB1501), mouse anti-TUJ1 (neuron-specific class III β-tubulin) (1:500 dilution) (BioLegend, catalog no. 801201), and Calbindin D28K (1:2000 dilution) (Swant antibodies, Switzerland, catalog no: CB38).

### Tissue processing of sarkosyl-inoculated mice

After 3 (C57BL/6J (n=6), *MAPT^0/0^* (n=3), ZH3-*Prnp^0/0^* (n=5), GPI^−^ *Prnp* (n=5)) or 6 (C57BL/6J (n=5), ZH3-*Prnp^0/0^*(n=3), GPI^−^ *Prnp* (n=6)) months post injection, mice were processed. After deep anesthesia with isoflurane, they were transcardially perfused with phosphate-buffered 4% PFA (pH 7,3) using a peristaltic infusion pump. The brains were dissected and postfixed overnight in the same fixative. As an internal control, sections from the frontal cortex (area 8/9) of a well-characterized AD sample (Braak stage VI) were processed in parallel. Next, postfixed brains were rinsed in 0.1M PBS and briefly in deionized water, and then stored in 70% ethanol at 4°C until paraffin inclusion. Following paraffin embedding, 10 µm coronal sections were obtained and mounted on gelatinized glass slides. For immunohistochemistry, paraffin tissue sections were deparaffinized in xylene for 20 min and rehydrated through a series of ethanol solutions followed by rinses in H_2_O. Selected de-waxed brain paraffin sections containing dorsal hippocampus were treated with DakoTarget retrieval solution (pH 9) (Dako, Denmark) at 95° in a Dako PT Link to retrieve protein antigenicity. After rinsing with 0.1M Tris-HCl pH 7.6, sections were rinsed in 0.1M PBS, and endogenous peroxidase activity was blocked by incubation in 2% H_2_O_2_ and 10% methanol dissolved in 0.1M PBS. After extensive rinsing, sections were incubated in 0.1M PBS containing 0.2% gelatin, 10% fetal bovine serum (FBS), 0.2% glycine, and 0.1% Triton X-100 for 1 h at room temperature. Afterwards, the sections were incubated with primary antibodies (all primary antibodies were diluted in 5% FBS, 0.1% Triton X-100, and 0.02% sodium azide in 0.1M PBS) overnight at 4°C. For bright-field visualization, tissue sections were then rinsed in PBS, and incubated for 2 h at room temperature with species-specific biotinylated secondary antibody (1:200 dilution) (Vector Laboratories) in 0.1M PBS. To reveal immunoperoxidase labeling, sections were incubated with an avidin-biotin peroxidase complex (ABC) kit following the manufacturer’s instructions (Vector Laboratories). Peroxidase activity was developed with 0.03% 3-3’-diaminobenzidine (DAB) and 0.01% H_2_O_2_. For fluorescence staining, sections were rinsed in 0.1M PBS and incubated with species-specific secondary antibodies, Alexa Fluor 488 or 568 (1:300 dilution) (Life Technologies), for 2 hours at room temperature. Finally, sections were incubated with 1 µg/ml Hoechst 33342 (ThermoFischer Scientific) diluted in 0.1M PBS for 10 min at room temperature, rinsed with 0.1M PBS, and mounted in Mowiol^TM^. Samples were photodocumented using an Olympus BX61 microscope equipped with a cooled digital DP72L camera and alternatively, fluorescence-stained sections were analyzed using a Zeiss Confocal microscope (Zeiss LSCM800).

### Tissue processing of P301S, P301S *Prnp^0/0^* and AAV-inoculated mice

Mice were anaesthetized at 9 months of age P301S (n =6), P301S *Prnp^0/0^*(n= 6) or 3 months after AAV infection (wild-type n = 4 ; ZH3 *Prnp^0/0^*n = 4) with isofluorane and perfused with 4 % PFA, cryoprotected in sucrose, frozen in dry ice and sectioned in a cryostat (40 μm thick). Free-floating sections were rinsed in 0.1 M PBS, and endogenous peroxidase activity was blocked by incubation in 3% H_2_O_2_ and 10% methanol dissolved in 0.1 M PBS. After rinsing, sections were incubated in 0.1 M PBS containing 0.2% gelatin, 10% normal horse serum, 0.2% glycine, and 0.2% Triton X-100 for 1 h at room temperature. Afterwards, the sections were incubated for 24 h at 4 °C with the primary antibody against AT8 (see below). After that, the sections were incubated with secondary biotinylated antibodies (2 h, 1:200 dilution) and streptavidin-horseradish peroxidase complex (2 h, 1:400 dilution). Peroxidase activity was revealed with 0.025% diaminobenzidine (DAB) and 0.003% hydrogen peroxide. Alternatively, primary binding antibody was detected using Alexa 488 tagged secondary antibody as above. After rinsing, the DAB and fluorescence processed sections were mounted onto slides as mentioned above. For quantification of AT8-positive neurons in the hippocampus (2–3 sections per mouse), the total number of positive cells (in 250 μm length of CA1) was counted using a ×40 objective (oil immersion Zeiss, N.A. 0.85). For immunofluorescence, AT8 immunostaining in the hilus (delineated using ImageJ^TM^ in 2–3 sections per mouse) was measured using the corrected total cell fluorescence (CTCF) method [19, 91, 100]. In the CTCF quantification, the granule cell layer and the labelling of the pyramidal layer of the CA4 were omitted.

### Primary neuronal cultures

For primary cortical cultures, E15.5-16.5 CD1 mouse (Charles River, France) embryo brains were dissected and washed in ice-cold 0.1M PBS containing 6.5 mg/ml glucose. The meninges were removed, and the cortical lobes were isolated. Tissue pieces were cut using a tissue chopper and trypsinized for 15 minutes at 37°C. After the addition of horse serum and centrifugation, cells were dissociated by trituration with 0.025% DNase (all from Sigma-Aldrich) diluted in 0.1M PBS with a polished glass pipette. Dissociated cells were plated at ∼3,000 cells/mm^2^ on plates (Nunc) coated with poly-D-lysine (Sigma-Aldrich). The culture medium was Neurobasal^TM^ supplemented with 2 mM glutamine, 6.5 mg/ml glucose, antibiotics (Penicillin and Streptomycin), 5% horse serum, and B27 (Thermo Fisher Scientific). After 72 hours, 5 µM Cytosine β-D-arabinofuranosidehydrochloride (AraC) (Sigma-Aldrich) was added for 48 hours to reduce the growth of dividing non-neuronal cells. Culture media was changed every two days. Some cultures were infected with AAV2r.*Syn*.P301L and treated 3 days post-infection with *P301S^+/−^* or sAD-derived sarkosyl extracts for 7 days. Next, cultures were fixed with buffered Methanol to remove soluble tau and immunostained using the human-specific HT7 antibody against Tau and β3-tubulin (TUJ1).

### Microfluidic devices

Microfluidic devices (MFD) were used in an optimized modification of our previous design of large dual-chamber, open neuronal co-culture, and of designs reported by [124]. The open microfluidic device consists of two main open chambers interconnected by 100 microchannels 950 μm in length, as used in previous studies (i.e., [128]). The large chamber areas (9 mm × 16 mm) facilitate effective cell culture and easy handling. The small cross-section areas of microchannels (10 µm *w* x 10 µm *h*) restrict the crossing of cortical neurons and dendrites but only permit the passage of neuronal axons. The microfluidic device was made of poly(dimethylsiloxane) (PDMS) using standard photolithography and soft lithography. For activity assays, primary cortical neurons from the entorhinal cortex (EC) and hippocampus (Hip) were obtained from E16.5 mice as above (wild-type = 10). Each microfluidic chamber was precoated with poly-D-lysine and seeded with 300,000 cells (EC or Hip neurons). Incubation media was as indicated above. Culture media was changed every two-three days.

### AAV infection of primary cultures in MFD with GCaMP6s, and calcium analysis of spontaneous neuronal activity

Spontaneous neuronal activity was analyzed in primary cortical cultures (wild-type = 5) after ptau and tau treatments. To allow the study of spontaneous neuronal activity, cultures in both chambers were infected with the genetically encoded calcium indicator (GECI) (AAV9.Syn.GCaMP6s.WPRE.SV40, Addgene) after 4 DIV. 2 x 10^7^ viral particles were used in the AAV infections. After 72 h post-infection, viral transduction was stopped by changing the culture medium, and at 8 and 11 DIV cultures were treated only in the EC compartment with 100nM or 200nM of recombinant tau (n = 5 MFDs) or ptau (n =6 MFDs) in 0.1M PBS. Medium was maintained from 8 to 11 DIV, removed at 11DIV and treated again with ptau and tau for two hours before recordings, no additional treatment were developed. Calcium time-lapse images were obtained in each of the devices at 8 DIV 2 hours after treatment (T0); 11 DIV (T3) and 15 DIV (T7). For the calcium imaging setup, we used an inverted Olympus IX71 fluorescence microscope equipped with a Hamamatsu ORCA Flash 4.0 camera, an LED light source (CoolLED’s pE-300white, Delta Optics, Spain), and an incubator from OkoLab (Izasa, Spain). Images of 16-bits were acquired using the camera software Olympus cellSens^TM^ (Olympus Corporation) at 500 ms exposure/frame for 5-6 min. The image size was 1024 x 1024 pixels, utilizing a 10x objective (Olympus UMPLanF). The videos were stored as muti-tiff files for further processing. Calcium traces were first analyzed using the NETCAL^TM^ software [99]. Identified neurons were associated manually in a homemade graphic using interphase (GUI) developed in Python 3 with a single region of interest (ROI). Each ROI includes only the perikaryon of the labelled neuron. Only cultures with dispersed non-grouped neurons were used. The number of selected ROIs in the study was 330 ± 69 (mean ± S.D.) for each video. The average fluorescence *Fi (t)* in each ROI *i* along the recording was then extracted, corrected for global drifts and artifacts, and finally normalized as *(Fi (t) — F _(0,i)_) / F _(0,i)_ = fi (t)*, where *F _0,i_* is the background fluorescence of the ROI. The time series of *fi (t)* was analyzed to determine sharp calcium transients. To reveal neuronal activity, normalized data files were processed using the protocols and algorithms developed by Sun and Südhof [121] using the algorithms in Matlab^TM^ software R2022b (lic. 48811179). This approach allowed us to analyze the overall spontaneous network activity as well as single-neuron dynamics. For biochemistry, microfluidic chambers were treated with tau (n = 8), ptau (n = 8), or 0.1M PBS (n = 8) as above at 7DIV and 4 additional devices were treated with total brain extract of P301S^+/−^ (as controls). Protein samples of cultured neurons (EC and Hip) were obtained 1, 3, and 7 days after treatment. In addition, the culture media of each reservoir was also simultaneously collected. Protein samples from cells and media were processed for Tau13, PHF1, and actin by western blot as indicated above.

### Statistical Analysis

Data in this manuscript are expressed as mean ± s.e.m. of at least four independent experiments (i.e., calcium analysis) unless specified. Means were compared using the Mann-Whitney *U* non-parametric or Turkeýs multiple comparison tests. The asterisks ∗, ∗∗, ∗∗∗ and ∗∗∗∗ indicate *P* < 0.05; P < 0.01; *P* < 0.001; and *P* < 0.0001, respectively. Statistical tests and graphical representations were performed with GraphPad Prism 10.0.1.218 (ID: 97F7835C03E).

## Results

### Characterization of the tau species used in the study

Western blotting of the sarkosyl-insoluble fraction of the AD patient (see Material and Methods) processed with anti-ptau Ser422 (pSer422) antibody revealed three bands of around 68, 64 and 60 kDa, together with an upper band of 73 kDa, several bands of about 50 kDa and some bands between 30-40 kDa, and two lower bands of truncated tau at the C-terminal, one of which was of about 25 kDa, as well as a smear of higher molecular weight represented large tau aggregates (Figure 1A). TEM of the same sarkosyl-insoluble fraction revealed the presence of typical paired helical filaments of tau (Figure 1B). In parallel, we analyzed by TEM the aggregation of the tau K18 fragment in the presence (Hep +) or absence (Hep -) of Heparin, to be further used as controls in our *in vitro* experiments (Figure 1C-D). As expected K18 only forms fibers in the presence of Heparin as revealed in TEM (Figure 1C-D). Next, we explored the seeding properties of the sarkosyl-insoluble fraction of the sAD patient using the Tau RD P301S Biosensor cell line. Thus, tau K18 (Hep+) (Figure 1F), sarkosyl-insoluble fractions of P301S transgenic mice (sP301S^+/−^) (Figure 1I) and the sarkosyl-insoluble fraction of the sAD patient (Figure 1J) were able to induce the formation of numerous eGFP aggregates in the cytoplasm of the cell line indicating their seeding properties in contrast to tau K18 (Hep -) (Figure 1G), the sarkosyl-insoluble fraction from P301S non-transgenic mice (sP301S^−/−^) (Figure 1H) and vehicle (Figure 1E).

**Figure 1.**
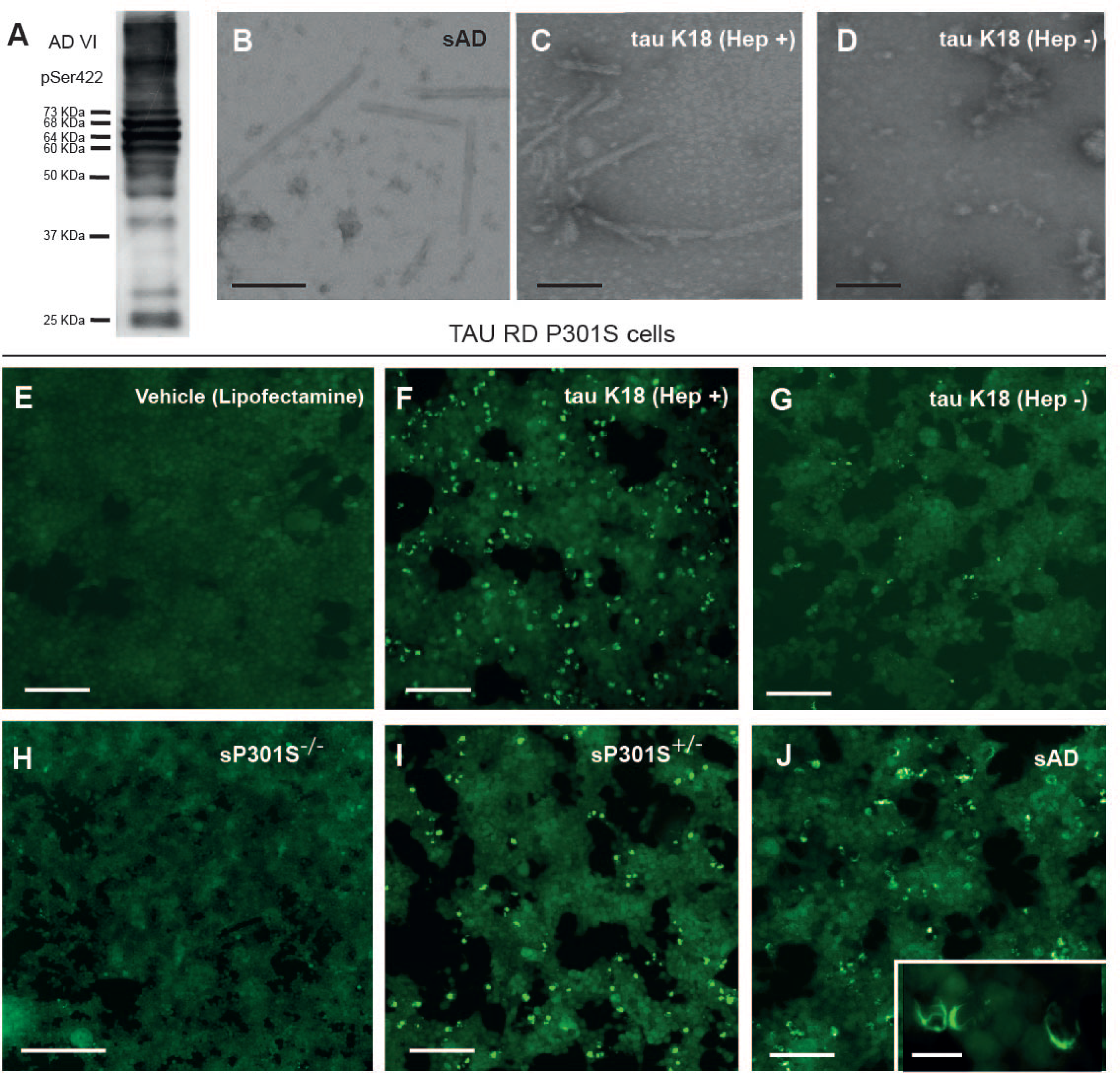
Characterization of the sarkosyl-insoluble fraction used in the study. **A)** Western blot of the selected sarkosyl-insoluble fraction from the AD patient (sAD) used in the manuscript immunoblotted using the pSer422 antibody. A discrete band pattern is observed from >70 kDa until 25 kDa with higher molecular forms. **B-C)** TEM negative staining pf the sAD (B) and tau K18 fragment after their fibrillation with (C) or without Heparin (D). **E-J)** seed competency experiments using the Tau RD P301S cell line of several samples including vehicle (E), fibrillar tau K18 (F), non-fibrillated tau K18 (G), sarkosyl-insoluble fraction from a P301S non-transgenic mouse (H), and P301S transgenic mouse (I) and sAD (J). Notice the presence of fluorescence aggregates in (F), (I) and (J) in contrast to vehicle (E), non-fibrillated tau K18 (G) and sP301S^−/−^. Scale bars: A-C: 0.5 μm. E-J = 200 μm; and the insert of J = 50 μm.

In additional controls, the treatment of the Tau RD P301S Biosensor cell line with i) monomeric tau Cy5 (Suppl. Figure 1A-C), ii) murine or human preformed fibrils of α-synuclein (Suppl. Figure 1D-E) or iii) sarkosyl-insoluble brain extracts from multiple system atrophy (MSA) (Suppl Figure 1F) or Parkinsońs disease (PD) (Suppl. Figure 1G) patients were unable to generate intracellular fluorescence aggregates. In contrast, treatments of the Tau RD P301S cells with sarkosyl-insoluble fractions from aging-related tau astrogliopathy (ARTAG, a 4R tauopathy) (Suppl. Figure 1H); Pick disease (PiD, a 3R tauopathy) (Suppl. Figure 1I) and globular glial tauopathy (GGT, a 4R tauopathy) (Suppl. Figure 1J) obtained in other studies [47, 49, 50] render intracellular fluorescence aggregates revealing tau seeding properties. In a second set of experiments, we isolated Triton X-100 tau insoluble fractions from Tau RD P301S cells treated during 24 h with sP301S^+/−^ (TIF-P+) and sP301S^−/−^ (TIF-P-). Next, freshly untreated Tau RD P301S cells were incubated with TIF-P+ and TIF-P-fractions as well as the vehicle-treated Tau RD P301S cells (TIF-V) (Suppl. Figure 2A-C). Results demonstrated that TIF-P+ fraction was able to generate *de novo* fluorescence aggregates in second passage experiments in Tau RD P301S cells (Suppl. Figure 2A) in contrast to TIF-P- and TIF-V fractions (Suppl. Figure 2B-C). Lastly, three P301S^+/−^ and two P301S^−/−^ 2-month-old mice (without endogenous ptau aggregation at this age) were inoculated with 2 μl of TIF-P+ extract in the dorsal hippocampus (Suppl. Figure 2D) and the presence of AT8-positive elements was analyzed 3 months later by immunohistochemistry (Suppl. Figure 2E-I). Revealed sections demonstrate the presence of AT8-positive labelling in the CA3 region hippocampus ipsilateral to the inoculation, the corpus callosum, and the contralateral hippocampus of TIF-P+ inoculated P301S^+/−^ (Suppl. Figure 2G, H-I)) mice but not in TIF-P+ inoculated P301S^−/−^ mice (Suppl Figure 2E,F). This demonstrates that the endogenous expression of a mutated form of tau enhances the aggregation of overexpressed tau under the presence of external Tau RD P301S-derived tau seeds. In parallel, similar results were obtained when primary mouse cortical neurons from wild-type mice were infected with AAV2r.*Syn*.P301L (Suppl. Figure 3) and treated with sarkosyl-insoluble extracts from P301S^+/−^ or P301S^−/−^ mice. Only seed-containing samples (P301S^+/−^) were able to induce the generation of HT7-positive neurons after AAV2r.*Syn*.P301L infection (Suppl. Figure 3D-F). However, treatment with the same sarkosyl-insoluble extract of AD to non-AAV2r.*Syn*.P301L infected cultures rendered negative results (at least at the time points analyzed) (Suppl. Figure 3I). In conclusion, in these experiments we characterized the sAD sample used in the study and, relevantly we demonstrate i) the specificity of the Tau RD P301S Biosensor cell line for tau seeds and ii) the seeding and spreading capacity of the TIF-P+ aggregates generated in the cell line in second passage experiments and after their *in vivo* inoculation in mutant tau P301S mouse model, and iii) the relevance of the endogenous expression of P301S/L in the appearance of human ptau aggregates.

### Characterization of PrP^C^ expression in mice strains used in the study

As indicated, the N-terminal and the CD domains of PrP^C^ have been described to bind to amyloids (see introduction for references). Thus, we aimed to analyze the pattern of expression of PrP^C^ in wild-type (*Prnp^+/+^*), ZH3 *Prnp^0/0,^* and GPI^−^ *Prnp* mice hippocampus by using western blot and immunohistochemical procedures (Figure 2) using the 6H4 anti-PrP^C^ antibody. Results (Figure 2A) revealed, as expected, the absence of PrP^C^ in ZH3 *Prnp^0/0^*mice extracts, and a band around 25 kDa in GPI^−^ *Prnp* brain extracts, as previously reported by [31], compared to the well-known three bands observed in wild-type derived brain samples (Figure 2A). Next, sections from wild-type (Figure 2B), ZH3 *Prnp^0/0^*(Figure 2C), and GPI ^−^ *Prnp* (Figure 2D) mice were processed for PrP^C^ immunostaining using the DakoTarget retrieval protocol (see Material and methods for details). Results demonstrate that PrP^C^ immunostaining in *Prnp^+/+^* was intense in all hippocampal layers being relevant in the CA1 except in the pyramidal layer and lower presence in the CA3 and dentate gyrus (DG) (Figure 2B). In contrast, ZH3 *Prnp^0/0^* (Figure 2C) and GPI^−^ *Prnp* (Figure 2D) hippocampus showed a pale uniform 6H4 labelling that was further corroborated in hippocampal sections from time-matched *Nestin*-*cre Prnp^flox/flox^*mice (Figure 2H). However, the gross anatomical cytoarchitecture of the hippocampus was maintained in all three genotypes as revealed by Calbindin immunostaining (Figure 2E-G).

**Figure 2.**
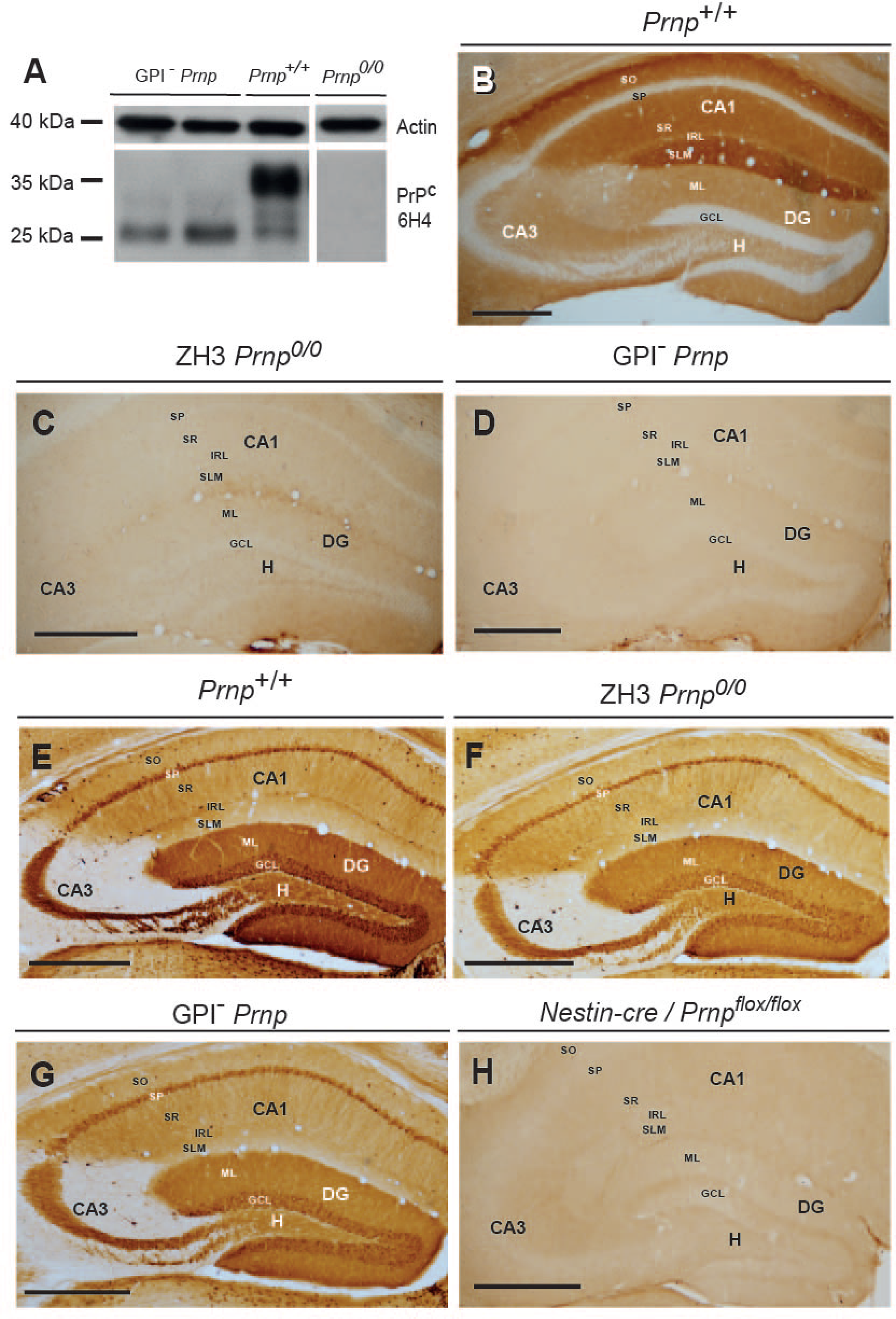
Expression of PrP^C^ and Calbindin in *Prnp^+/+^*; ZH3 *Prnp^0/0^* and GPI^−^ *Prnp.* **A)** Western blot of brain extracts from the three genotypes immunoblotted using actin and the 6H4 against PrP^C^. Notice the absence of PrP^C^ labeling in ZH3 *Prnp^0/0^*, the well described triple band in the wid-type mice and the lower band described in the anchorless PrP^C^ mice. **B-D)** Low power photomicrographs illustrating coronal hippocampal sections immunoreacted against PrP^C^. After DAB development, immunolabelling was intense in *Prnp^+/+^* hippocampus especially in the CA1 region (B). In contrast a pale labeling was observed in ZH3 *Prnp^0/0^* and GPI^−^ *Prnp* hippocampi (C-D). **E-G)** Calbindin immunostaining in the three genotypes demonstrating that the laminar and the general cytoarchitectonics is maintained in all genotypes. **H)** 6H4 immunostaining in the hippocampus of *Nestin-cre* / *Prnp^flox/flox^* mouse. As observed, very few labelling can be observed after immunostaining. Abbreviations: CA1-CA3 = *cornus ammonis* 1 and 3; DG = Dentate gyrus; H = Hilus; SO = *stratum oriens;* SP = *stratum pyramidale;* SR = *stratum radiatum*; IRL = interphase *radiatum-lacunosum moleculare*; SLM = *stratum lacunosum-moleculare*; ML = molecular layer; GCL = granule cell later. Scale bars: B-H = 500 μm.

### Absence of PrP^C^ expression or the expression of anchorless PrP^C^ do not impair endogenous tau aggregation after sAD inoculation

As indicated, several studies point to a PrP^C^-tau interaction, but no data is reported concerning a putative role of the physiological levels of PrP^C^ in promoting ptau generation and propagation in sarkosyl-insoluble inoculated mice. Thus, 4-5 month-old ZH3 *Prnp^0/0^*, GPI^−^ *Prnp,* and wild-type mice were inoculated with the same amount of sAD in the cortex/corpus callosum and hippocampus and analyzed 3- and 6 months later (Figure 3). Results indicate the presence of numerous AT8-positive deposits in inoculated mice irrespective of their genotype and post-inoculation time (Figure 3). Controls lack the presence of AT8-positive deposits. No clear differences in the ptau deposits and its cellular presence were observed after the comparison of all *Prnp* genotypes with wild-type mice at the two time-points analyzed (Figure 3). AT8-positive inclusions of different morphologies (pre-tangles, threads, and coiled bodies but not neurofibrillary tangles) were observed mainly in neurons and oligodendrocytes immunolabeled with AT8 or MC-1 antibodies (Figure 3A-D), but not in astrocytes (Figure 3E), expanding from the injection site in the cortex through the cortical white matter from the ipsilateral to the contralateral hemisphere (Figures 3 and 4). Relevantly, in our experiments, numerous AT8-positive ptau deposits were observed in axonal tracts suggesting relevant axonopathy after cortical sAD inoculation in all analyzed genotypes (Figure 3F). These ptau inclusions were also positive for X34 (Figure 3G). Few neurons were observed labelled in the contralateral neocortex or hippocampus being mainly observed near the injection site (Figure 3H-I). Relevantly, all mice treated with AD-derived sarkosyl-insoluble fractions showed double labelled cells containing AT8-positive deposits and intense P62 labelling (see Suppl. Figure 4 for some examples) suggesting that treated cells modified proteostasis or activates a putative necroptosis process after the inoculation. However, and relevantly our data demonstrates that PrP^C^ absence does not impair ptau seeding in affected neurons and oligodendrocytes.

**Figure 3.**
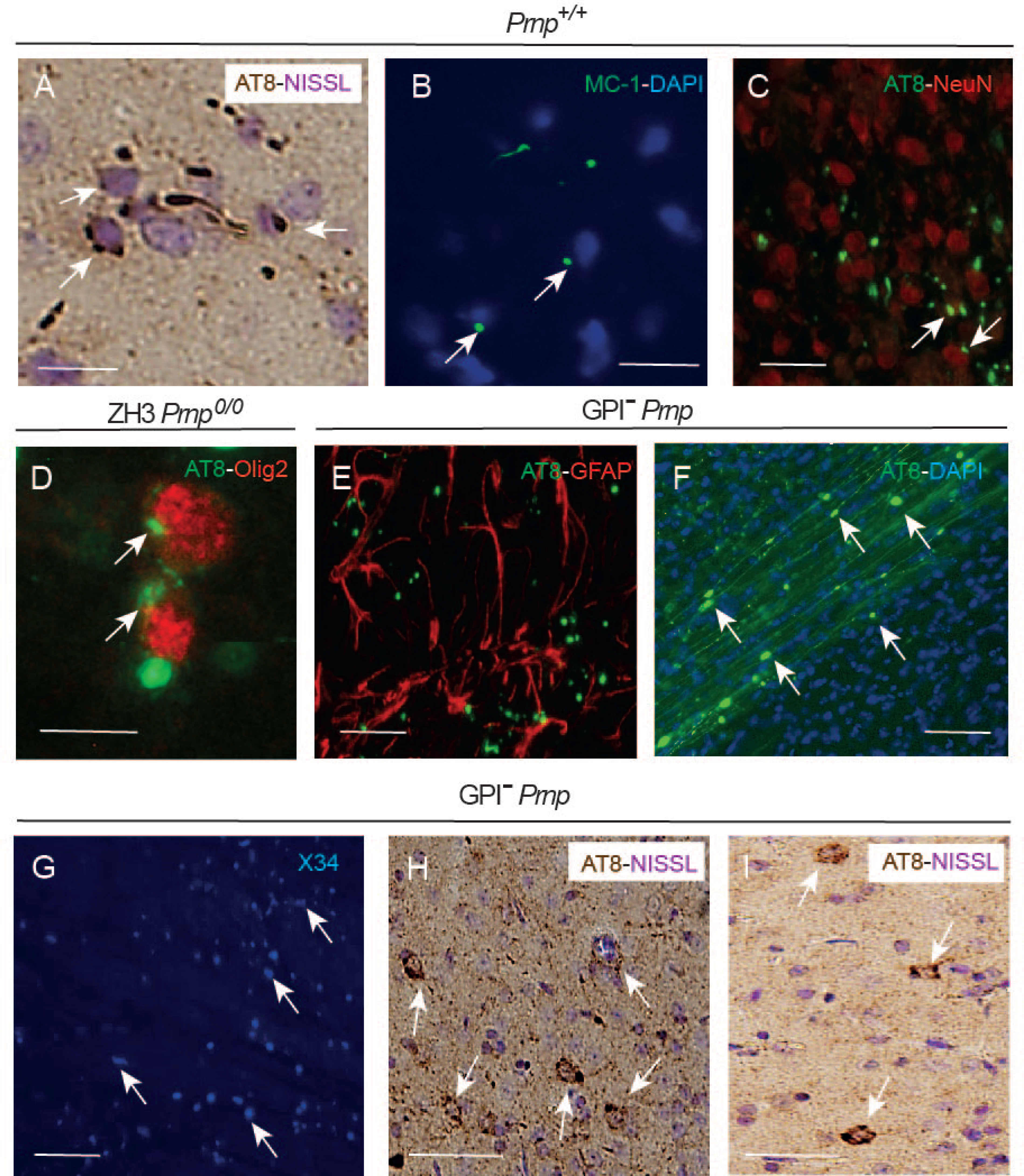
AT8- and MC1 ptau positive deposits in the different genotypes of the study. Examples of the different immunostainings developed after sAD inoculation at 3 (H-I) and 6 months post inoculation time (A-G). **A)** AT8-positive grains (arrows) and treads in the SLM of the CA1. The section was Nissl counterstained. **B)** MC-1 positive treads and grains (arrows) in the lower cortical layers after sAD inoculum. The section was counterstained with DAPI. **C)** Double immunofluorescence against NeuN (red) and AT8 (green) in lower cortical layers. Notice the presence of some graisn in NeuN positive neurons (arrows). **D)** Double immunofluorescence illustrating examples of Olig2– positive oligodendrocytes (red) containing AT8-positive inclusions (green and arrows) in the white matter. **E)** Absence of double labeled GFAP (red) and AT8 (green) in the CA1 of inoculated mouse. **F)** Immunofluorescence against AT8 (green) illustrating the relevant labelling of axonal tract in the white matter with large aggregates (arrows) indicating axonopathy. The section was counterstained with DAPI. **G)** Low power photomicrograph illustrating the presence of X34 positive labelling in the white mater of inoculated mice. In fact, the treatment with Sudan black prevented X34 immunostaining (not shown). **H-I)** Examples of AT8-positive neurons (arrows in H and I) in the retrosplenial cortex after sAD inoculation in the ipsilateral (H) or contralateral (I) cortex. The section was Nissl counterstained. Scale bars: A-D = 50 μm; E-G = 100 μm; H-I = 50μm. The genotype of the mouse is illustrated in each picture.

**Figure 4.**
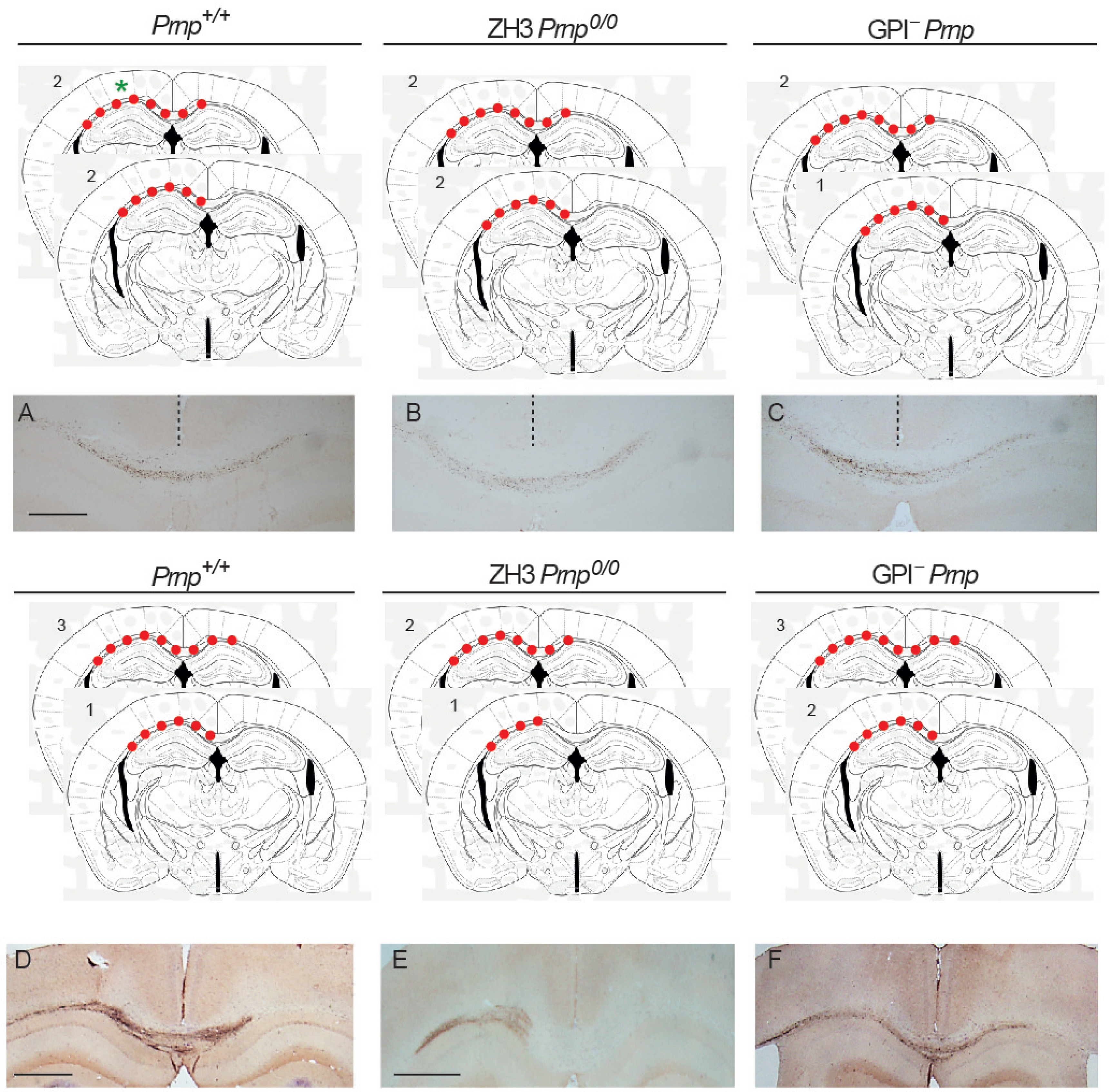
Propagation of AT8-positive deposits in the white matter between 3 and 6 months post sAD inoculation in the different genotypes. For each genotype 2 schemes are included. In each scheme the distribution of the AT8-positive deposits is illustrated with red circles. The inoculation region is illustrated by a green asterisk in (A), and the number of inoculated mice showing the distribution indicated in the scheme is also indicated in the upper left corner. **A-C)** Photomicrographs illustrating examples of the distribution of the AT8-positive deposits in the three different genotypes 3 months after sAD inoculation. **D-F)** Photomicrographs illustrating examples of the distribution of the AT8-positive deposits in the three different genotypes 6 months after sAD inoculation. The interhemispheric fissure is labelled with a dashed line in (A-C). Notice the increase in the overall labeling between 3 and 6 months post sAD inoculation. However, the distribution of the ptau deposits is similar between genotypes and post inoculation times. Scale bars: A = 250 μm pertains to B-C; D = 500 μm; E = 500 μm pertains to F.

After immunostaining at 3 and 6 months, the extension of AT8 positive ptau in the inoculation site and the white mater was analyzed (Figure 4). For wild-type 4 of the 6 inoculated mice showed AT8-positive deposits in the white matter reaching the interhemispheric middle line (Figure 4) and 2 of them crossed towards the contralateral hemisphere (Figure 4A). For ZH3 *Prnp^0/0^* and GPI^−^ *Prnp*, similar results were obtained, and in 50% of inoculated mice, the deposits reached the interhemispheric middle line 3 months post-inoculation (Figure 4B-C). At 6 months post-inoculation, AT8 immunostaining was intense compared to 3 months but very few changes were observed in the distribution of the AT8-positive deposits in the white matter and the inoculation site (Figure 4D-F). For wild-type, 3 of 5 mice showed AT8-immunoreactive deposits crossing the interhemispheric middle one reaching the interhemispheric middle (Figure 4D). For ZH3 *Prnp^0/0^* and GPI^−^ *Prnp* the labelling was similar to those observed 3 months after inoculation (i.e., 5 of 6 GPI^−^ *Prnp* mice showed labelling on the corpus callosum and contralateral hemisphere (Figure 4F)) and for ZH3 *Prnp^0/0^*2 mice showed contralateral labelling but one mouse showed absence of the AT8-positive deposits in the contralateral site (Figure 4E). In conclusion, results reported a similar seeding and propagation of AT8-positive deposits after inoculation of AD-derived sarkosyl-insoluble fraction between wild type, GPI^−^ *Prnp^0/0^* and ZH3 *Prnp^0/0^* mice between 3 and 6 months. Since only small differences can be observed between inoculated mice, current data suggests a minor role of PrP^C^ and the expression of the anchorless PrP^C^ in the propagation of ptau after sAD sarkosyl-insoluble inoculation *in vivo*.

### Pathologic AD sarkosyl-insoluble fraction inoculation triggers endogenous mouse ptau generation and exon 10 splicing

Next, we aimed to characterize in more detail the observed ptau aggregations taking into account that although some studies reported seeding in inoculated wild-type mice (i.e., [11, 33, 61]), most of the studies aiming to determine the seeding and spreading capabilities of different tau seeds were developed using transgenic models overexpressing mutated forms of human tau (i.e., P301S (i.e., [4, 18, 33]) or ALZ17 (i.e., [32]) or 4R human tau. (i.e., [47]). In addition, we were able to determine in previous experiments, that only when the treated neurons express a human mutated form of tau can they generate human tau shortly after exposure to human-derived tau seeds. Thus, we first inoculated *Mapt^0/0^* mice with the sAD inoculum. Results reported the absence of AT8-positive labelling in processed sections (Figure 5), demonstrating that ptau inclusions previously observed in ZH3 *Prnp^0/0^*, GPI^−^ *Prnp,* and wild type mice after 3- or 6-month post inoculation were of endogenous mouse origin. This was also corroborated by immunostaining using T49 (mouse-specific) and Tau13 (human-specific) anti-tau antibodies (Figure 5). Thua, immunolabelled sections demonstrated that pSer422-positive ptau aggregates were only labeled with the T49 mouse-specific anti-tau antibody. With some differences, this was also observed by [38] or [76]. Lastly, we aimed to determine whether the observed ptau inclusions contained 3R or 4Rtau isoforms by using specific antibodies (Figure 5). Immunoreacted sections demonstrated the presence of 3R and 4Rtau isoforms in pSer422-positive aggregates. These data demonstrate that, although PrP^C^ can interact and transport intracellularly with different forms of tau [25] or [35], PrP^C^ absence or presence in the extracellular space does not impair significantly the seeding properties of sAD. Lastly, the generated 3R or 4R pSer422tau deposits are of mouse origin and showed low spreading properties to contralateral neurons, being localized in ptau-positive neurons and a low number in oligodendrocytes in the cortical white matter.

**Figure 5.**
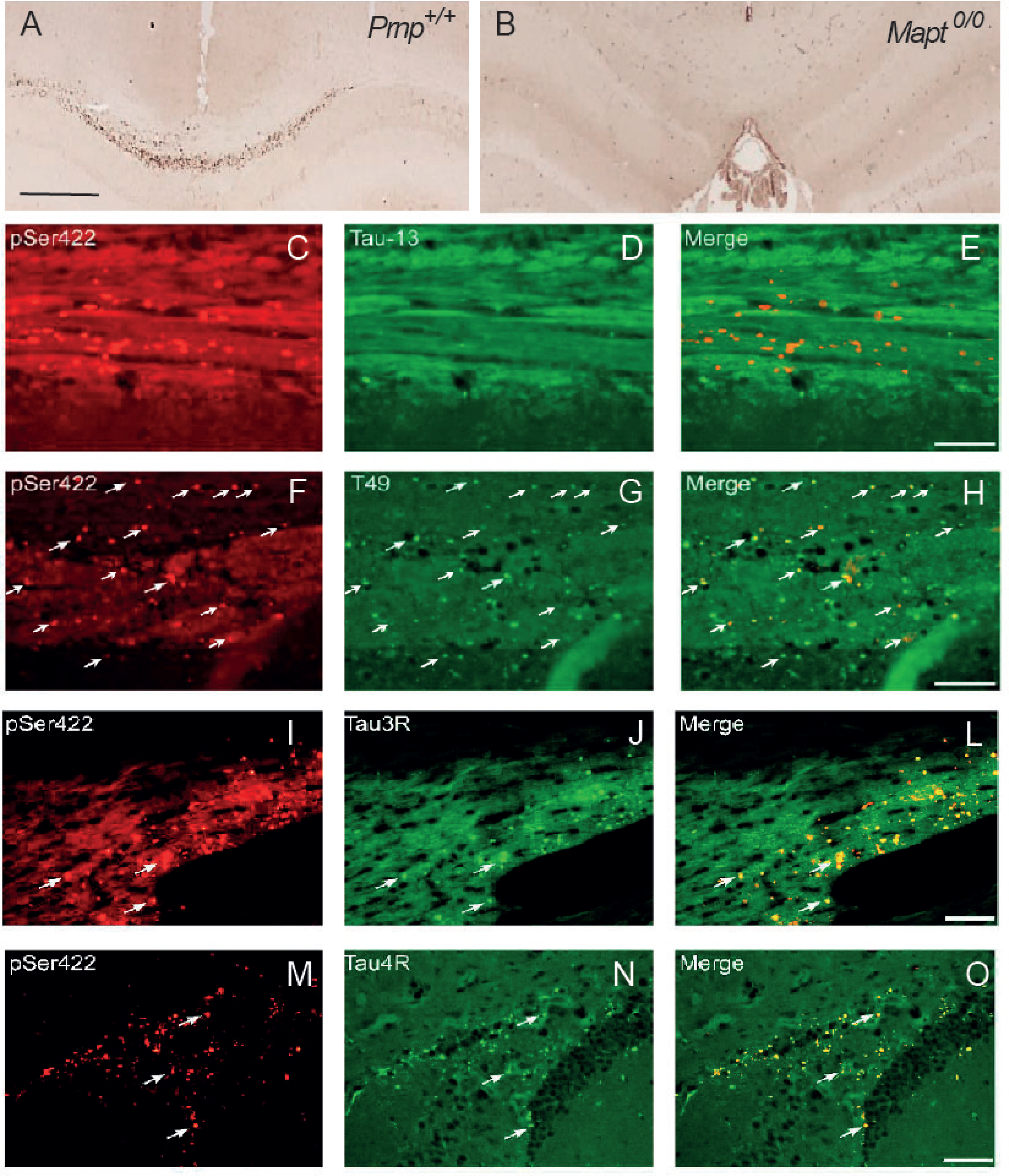
Characterization of the ptau deposits observed in the different genotypes. **A)** This is the same example showed in Figure 4A of a wild-type after sAD inoculation and immunoreacted using AT8 antibody. **B)** Low power photomicrograph illustrating the interhemispheric fissure of a *Mapt^0/0^* mouse after sAD inoculation and immunostating against ptau (AT8 antibody). Notice the absence of AT8-positive elements in deficiency of endogenous mouse tau. **C-E)** Double immunofluorescence images illustrating pSer422 (C) and Tau 13 (D) immunostaining. Notice the absence of double labelled deposits. **F-H)** Double immunofluorescence images illustrating pSer422 (F) and Tau 49 (G) immunostaining. Notice the presence of double labelled deposits (arrows in F-H) demonstrating tha the observed ptau is of mouse origin. **I-O)** Double immunofluorescence images using pSer422 (I and M) and Tau 3R (J) or Tau 4R (N) specific antibodies. Notice the presence of double labeled deposits in L and O indicating that sAD inoculum can modulate exon 10 of the *Mapt* gene. Scale bars: A = 500 μm pertains to B, E, H, L and O = 100 mm pertains to C-D, F-G, I-J and M-N respectively.

### Human full-length tau or ptau treatments enhanced spontaneous neuronal activity in a dose-dependent manner of primary mouse neuronal cultures

After the above-observed results, we aimed to determine whether both ptau and tau can be uptaken by treated EC neurons and propagated to untreated Hip neurons in MFDs (after 1, 3, and 7 days post-treatment). Our two chamber MFDs with 950 μm microchannels showed clear microfluidic isolation (see [128] for details)(Figure 6) in contrast to other studies [107]. In the experiments, 10 μg of a total brain extract from a P301S^+/−^ mouse of 11 months of age was also used as a control. In addition, changes in the activity of EC neurons were analyzed as indicated above in longitudinal experiments (Figures 7). Western blot analysis of ptau and tau showed the appearance of Tau13 in Hip neurons at 7 days after treatment with ptau in the EC (Figure 6B). In addition, ptau, tau, and P301S extract can be detected in Hip neurons but not in the culture media after 7 days post-treatment (Figure 6C). However, Tau13 detected in Hip neurons after treatment was not labelled using PHF1 antibody (Figure 6D). In parallel, calcium analysis of spontaneous neuronal activity using AAV9.*Syn*.GCaMP6s reported a very low number of synchronous events at T0 irrespective of the treatment with vehicle, tau (100 nM), or ptau (100 nM) (vehicle = 1; tau = 1.3 ± 0.33 or ptau =1) (Figure 7). Number of synchronous events increased at T3 and T7, but more relevantly after ptau or tau treatments (100 nM) (T3 = vehicle: 10.50 ± 0.8; tau: 13.20 ± 2.76; ptau: 11.0 ± 1.51; T7 = vehicle: 7.5 ± 0.5; tau: 16.33 ± 1.803; ptau: 9,01 ± 1.55) (Figure 7). This correlates with an increase in the number of events per neurons/time at the different stages (vehicle: T0 = 0.08 ± 0.007, T3 = 0.369 ± 0.05, T7 = 1.798 ± 0.14; tau: T0 = 0.57 ± 0.14; T3 = 2.043 ± 0.30; T7 = 3.183 ± 0.173; ptau: T0 = 0.10 ± 0.04, T3 = 0.33 ± 0.11, T7 = 1.065 ± 0.44). The main statistics values can be seen in Figure 7 and all values in Suppl. Table 1. Surprisingly, the number of synchronous events largely increased (≈ 8-10 times) from T0 when ptau or tau were added at 200 nM (T0 = tau: 11.50 ± 1.5; ptau: 8,0 ± 3). A relevant decrease in the synchronous and the general activity of the cultures was observed shortly at T3 (T3 = tau: 3 ± 0.1; ptau:1) and all 200 nM treated cultures died before T7 in contrast to the 100 nM tau or ptau treatments (Figure 7). Thus, present results demonstrate that exogenous treatment with either fibrillar ptau or tau increased spontaneous neuronal activity but reduces neuronal activity in a dose-dependent manner, leading to premature cell death at higher concentrations.

**Figure 6.**
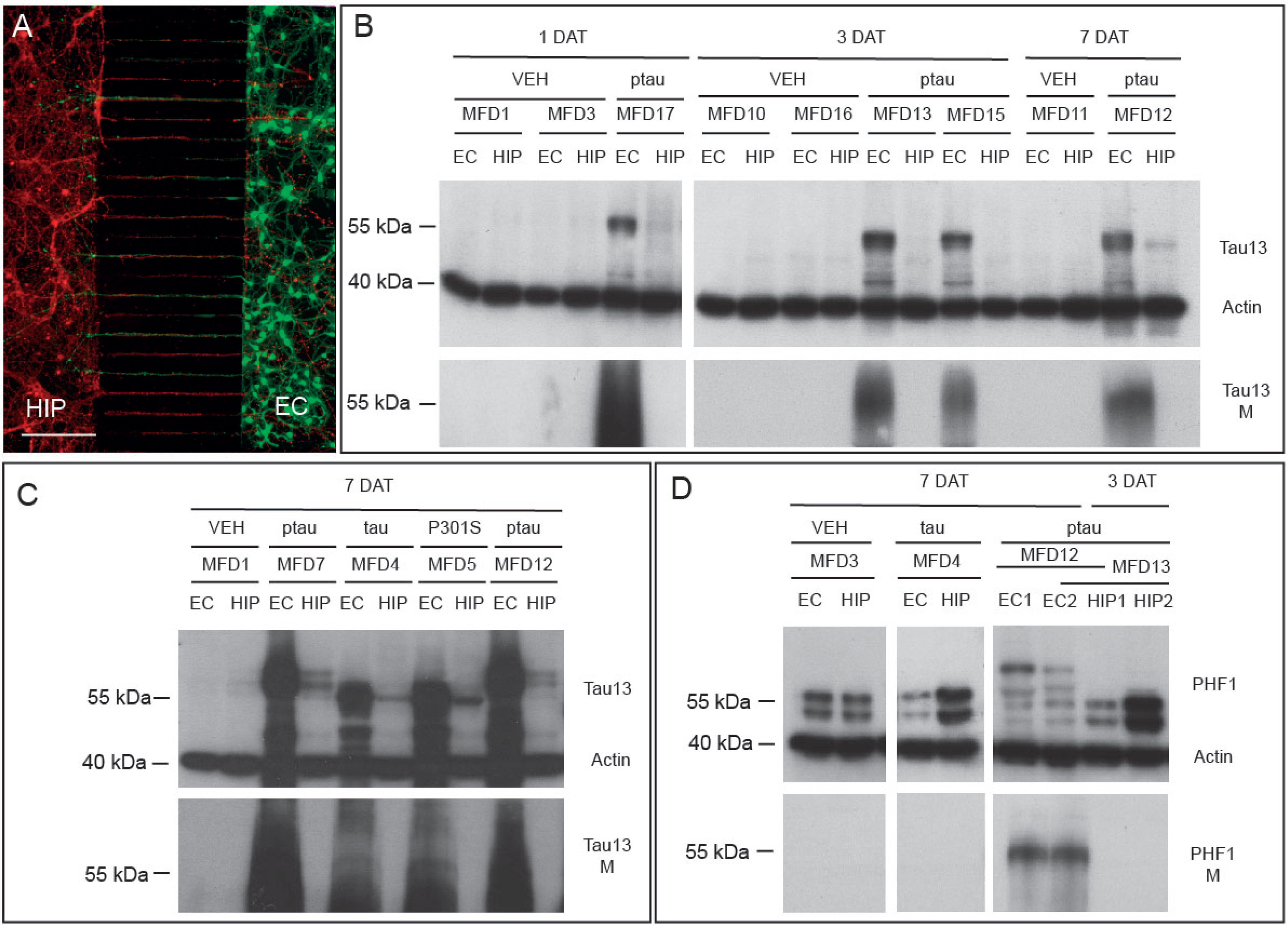
Uptake and transport between EC and HIP neurons of different forms of tau in MFDs. **A)** Example of a MFD used in the experiments of Figure 6 and 7. EC and Hip were seed in the different chambers, and 7 days after seeding EC neurons were labelled with AAV9.*Syn*.GCaMP6s and HIP neurons with AAV9.*Syn*.JRCaMP1b. As observed neurons interconnect by crossing their axons though the microchannels of 950 μm length. **B-D)** Western blots of treated MFDs with vehicle (VEH), fibrillar tau, fibrillar ptau (B-D) and soluble brain extract from P301S mouse (C). The days after treatment (DAT) are indicated in each panel. For each MFD, the result of the western blot in the cells (up) and the culture media (down, i.e., Tau13 M in B and C) is illustrated. In addition, the number of the MFD used is included and the bottom line of each MFD links the EC and HIP chambers of the device. Cells and media extract were immunoblotted using Tau13 and actin (B-C) or with PHF1 and actin (D). As observed in (B), labelling of Tau13 appeared in the HIP cells after ptau incubation of EC chamber only after 7 DAT. The same happens for others tau species (C). However, human tau transported between cells become de phosphorylated in recipient cells (D). Notice that in all cases the observed transport in a cell mediated transport and no diffusion in the media of the tau species can be detected. Scale bar: A = 450 μm.

**Figure 7.**
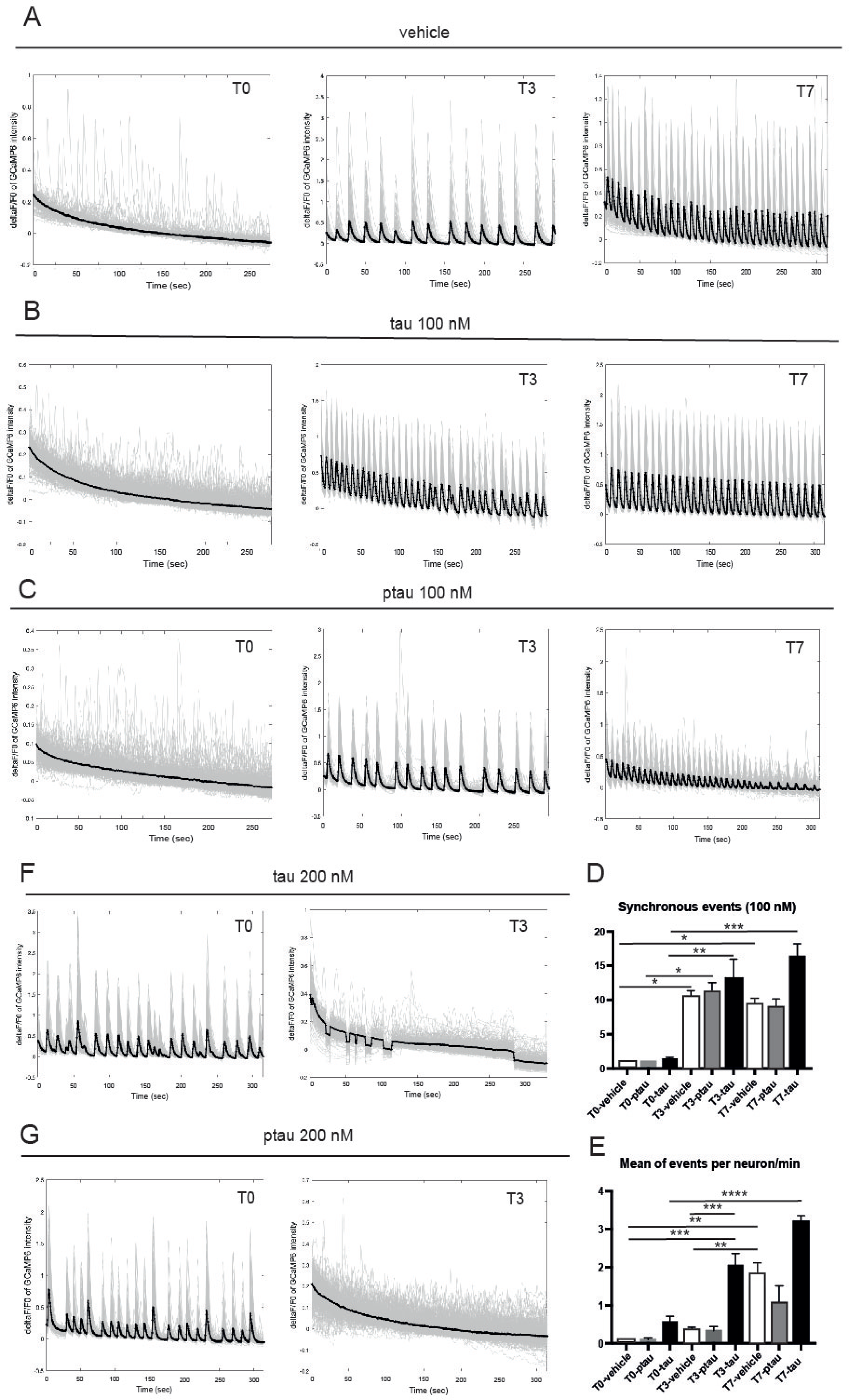
Analysis of the spontaneous neuronal activity in GCaMP6s expressing neurons of the EC after treatments with ptau or tau at different concentrations. **A-E)** Representative ΔF/F0 plots of GCaMP6s-expressing EC neurons generated after the changes in fluorescence at T0, T3 and T7 in the EC neurons. In each graph the changes in the fluorescence levels of each neuron corrected to the background is plotted in grey and the mean of the observed calcium traces is plotted as the dark line. Synchronous events can be easily identified. As indicated, cultures were treated in the EC with vehicle (0.1 PBS)(A); tau 100 nM (B); ptau 100 nM (C), tau 200 nM (D) or ptau 200 nM (E). Notice the increase of the synchronous events between T0 and T7 in (A-C). After 200 nM incubation neurons showed a increased spontaneous activity at T0 (D-E) being decreased at T3. These cultures died before T7. **F)** Graph illustrating the number of synchronous events in the different time points and treatments (100 nM). **G)** Graph illustrating the frequency as the mean of events per neurons and min in the different conditions. Asterisks indicate statistical significance: * *P* < 0.05; ** *P* < 0.01; *** *P* < 0.001; Turkeýs multiple comparison test. Main statistical differences are illustrated but all the statistical values can be seen in Suppl. Table 1.

### Decreased AT8 positive neurons in the hippocampus of P301S-ZH3 *Prnp^(0/0)^*mice compared to P301S at 9 months of age

As previous results point out that the absence of PrP^C^ does not play a crucial role in reducing the seeding of sAD fractions and few effects can be seen in propagation, we generated a double mutant mouse lacking *Prnp* in the P301S transgenic mice (Figure 8A-B). As reported, these mice start to generate ptau aggregates after 6 months of age [133]. At 9 months of age, coronal brain sections were fixed and processed for AT8 immunoreactivity and the number of AT8-positive cells was counted in the pyramidal layer of the CA1. Results indicated the presence of 15.03 ± 0.566 and 11.04 ± 0.68 (all mean ± s.e.m.) AT8-positive neurons in P301S and P301-ZH3 *Prnp^0/0^* mice respectively (Figure 8C), demonstrating that the genetic ablation of *Prnp* reduces the number of AT8-positive CA1 neurons in P301S^+/−^ mice. In parallel, 3-month-old wild type, *Mapt^0/0^*and ZH3 *Prnp^0/0^* mice were infected with AAV2r.*Syn*.P301L in the dentate gyrus and processed 3 months post-infection with AT8 antibodies (Figure 8D-G). After immunochemistry, the AT8-positive fluorescence observed in the hilus was analyzed by CTCF. Obtained values were statistically non-significant (wild-type: 1.278 ± 0.21; *Mapt^0/0^*: 1.592 ± 0.32 and ZH3 *Prnp^0/0^*: 0.76 ± 0.181. all mean ± s.e.m.) but lower values were observed in the ZH3 *Prnp^0/0^* genotype (*P* = 0.44 wild-type *vs* ZH3 *Prnp^0/0^* and *P* = 0.27 *Mapt^0/0^ vs* ZH3 *Prnp^0/0^*, One way ANOVA, Mann-Whitney *U* non-parametric test) (Figure 8G). In conclusion, the absence of *Prnp* decreases the presence of AT8-positive neurons in the P301S^+/−^ mice hippocampus.

**Figure 8.**
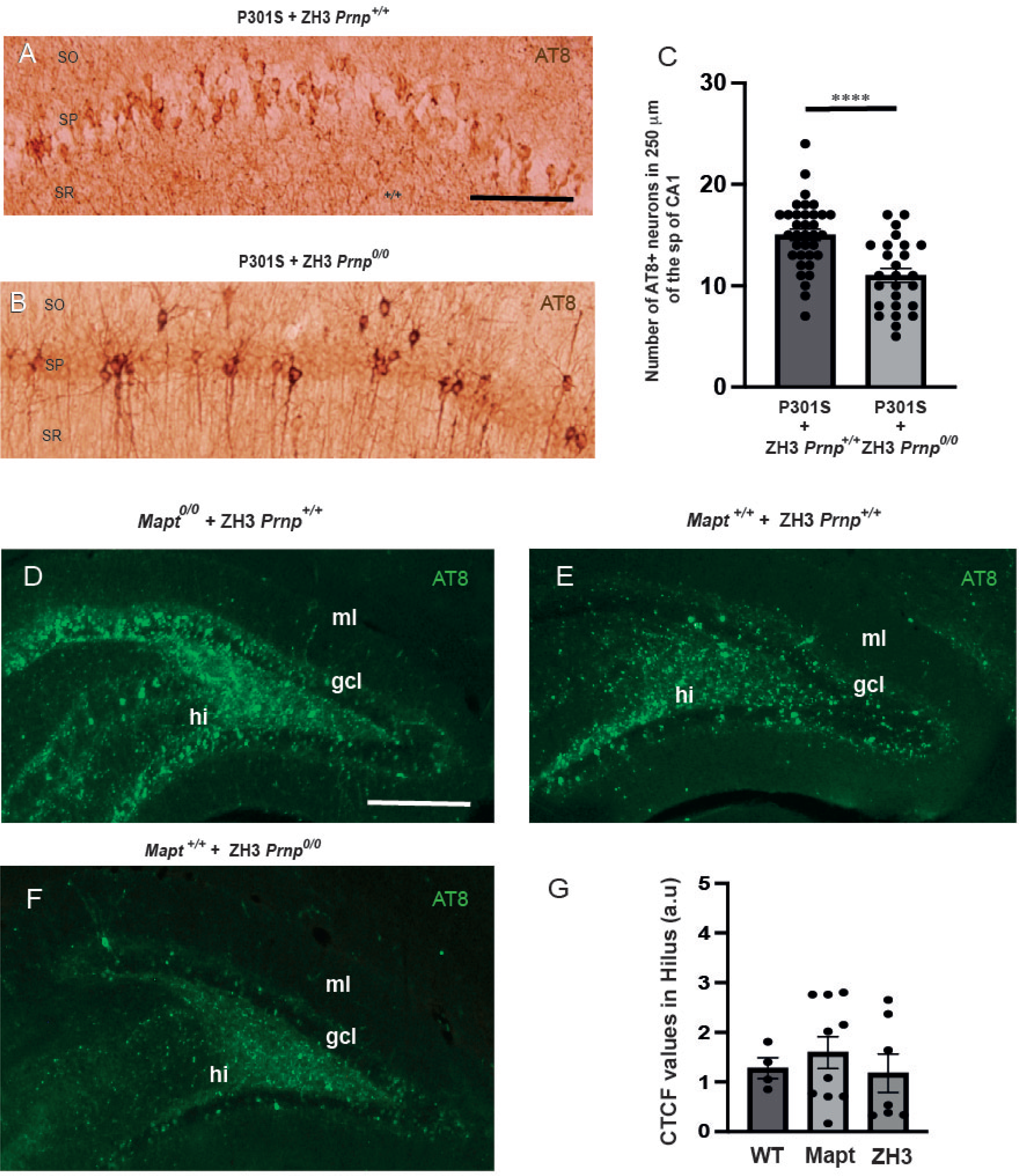
Effect of the *Prnp* absence in AT8-positive immunoreactivity in P301S mice. **A-B)** Photomicrographs illustrating the AT8-positive neurons in the pyramidal layer of the CA1 in P301S + ZH3 *Prnp^+/+^* (A) and P301S + ZH3 *Prnp^0/0^*(B) hippocampus. **C)** Histograms illustrating the quantification of the immunoreacted section showed in (A-B). Each dot represents a quantified value. **D-F)** Low power fluorescence photomicrographs illustrating examples of AT8-immunoreacted dentate gyrus of *Mapt^0/0^* + ZH3 *Prnp^+/+^* (D), *Mapt^+/+^* + ZH3 *Prnp^+/+^*(E) and *Mapt^+/+^* + ZH3 *Prnp^0/0^* after the infection with AAV2r.*Syn*.P301L virus (see Material and Methods for details). **G)** CTCF quantification in the hilus of immunoreacted sections as in (D-F). For plotting purposes, x-axes nomenclature was WT for wild-type, Mapt for *Mapt^0/0^* + ZH3 *Prnp^+/+^* and ZH3 for *Mapt^+/+^* + ZH3 *Prnp^0/0^*. Each dot represents a quantified value. Abbreviations as in Figure 2. Scale bars: A = 100 μm pertains to B; D = 250 μm pertains to E-F. Asterisk indicate statistical significance (****, *P* < 0.0001; Mann Whitney *U* non-parametric test)

## Discussion

PrP^C^ is involved in a plethora of functions during the development and in the adult CNS (i.e., [57, 87, 129]). In addition, it plays roles during neurodegeneration and other neural disorders (i.e., [23, 42, 94, 111, 131, 135, 136]). During neurodegeneration, PrP^C^ interacts with several amyloids mainly by its CD or N-terminal domains (i.e., [79, 112, 115, 127, 138] as suggested some years ago [105]. From a physiological point of view, oligomeric amyloids modulate synaptic activity (i.e., [62] being in some studies considered as neuroprotective in heathy non-pathological situations (i.e., [16, 120]. In fact, the physical interaction of PrP^C^ with, mainly oligomers of Aβ, α-synuclein, tau, and TDP43 has been already demonstrated (i.e., [55, 114, 115, 125, 128]). Indeed, increasing PrP^C^ levels by transfection in supraphysiological levels that normally do not occur *in vivo*, enhanced tau uptake in overexpressing cells (i.e. [25, 107]. In a previous study, we and others demonstrated that PrP^C^ plays a relevant role in the propagation of sonicated α-synuclein fibrils. Here, their participation is minor in the seeding and spreading of AD sarkosyl-insoluble fractions. Indeed, our *in vivo* experiments reported no differences in seeding or spreading of sarkosyl-insoluble AD samples in the *Prnp* mice compared to controls in the study, leading to the notion that other factor/s different than PrP^C^ play more relevant roles during ptau seeding and spreading after sarkosyl-insoluble inoculation. Although no clear differences between genotypes were observed, the generated ptau deposits are of mouse origin affecting neurons and oligodendrocytes as also reported in some studies using wild type mice (i.e, [8, 11, 38, 76]). In addition, in our experiments, sAD inoculum was able to change exon 10 splicing of endogenous mouse *Mapt* gene. Tau deposits in all three genotypes inoculated with sAD sarkosyl-insoluble fractions are composed of 4R and 3R tau in contrast to [11]. This described that a change from fetal to adult tau isoform expression occurs in mice and humans [63] and can be observed during maturation in human embryonic or induced pluripotent stem cell cultures [68]. However, a shift in the expression of 4R to 3R tau has been reported in oligodendrocytes following cerebral occlusion in adult rats and mice (see [46] for review), thus suggesting exon 10 splicing modulation, under determinate unhealthy conditions (i.e. ischemia), could be detrimental. In fact, recently the 2R and 3R domains of tau have been determined crucial for tau aggregation [9, 86, 117].

In the study, the role of PrP^C^ in AT8-positive ptau generation was different between transgenic mutant mice and to a lesser extent in AAV infections when compared to sAD inoculated wild type mice. In several studies using sarkosyl-inoculated mice, shortly after the inoculation, most if not all ptau from human inoculum cannot be observed in the injection site (i.e., [8, 11, 61, 76]). Our microfluidic experiments reported that fibrillar forms of human tau (phosphorylated and unphosphorylated) can be uptaken and transported between primary cortical neurons (EC > Hip cells) shortly after 7 days *in vitro*. Indeed, human tau is present in recipient cells but in a non-phosphorylated form confirming the absence of human ptau after sAD inoculation in wild-type mice. On the other hand, the presence of a relevant amount of exogenous normal tau in mouse neurons might trigger neurotoxic effects such us axonopathy and neurite fragmentation *in vitro* [67, 81]. This was also determined by using lentiviral delivery of human wild-type tau in rats [22]. In addition, our calcium imaging analyses corroborate that incubation with fibrillar tau in primary neuronal cultures largely increased spontaneous neuronal activity and, in a dose-dependent manner, loss of synchronicity and cell death. In fact, absence of tau largely impair the synchronicity in cultured neurons [29], but relevantly neuronal activity increases tau delivery [132] as well as tau propagation and pathology [113, 130]. Moreover, a recent study points out that in a genetic model of tauopathy with progressive ptau aggregation, disrupted neuronal dynamics at the network level correlate with the progression of the tauopathy [92]. As indicated, all sAD-inoculated mice showed relevant axonopathy with few labelled cells after inoculation and poor cell-to-cell propagation of ptau. Similar results concerning the location of ptau deposits after sarkosyl-insoluble inoculation in wild-type mice were described in some studies by (i.e., [8, 32, 33]). In fact, it has been described that oligomeric species of tau can spread to different brain locations when compared to fibrillar aggregated forms (i.e., [76, 80]) since soluble tau species increase seeding when compared to large amyloid fibrillar forms [58, 80]. Relevantly, after sAD inculation numerous cells are double labelled with P62 and AT8 antibodies. P62 is a critical molecule participating in several processes of autophagic flux including the assembly of autophagosome, fusion of autophagosomes with lysosomes, and cargo degradation [71]. In fact, P62 is also degraded with them [85]. Indeed, p62 accumulation correlated with increasing ptau seed accumulation after inoculation in our study and in [17]. Although we cannot rule out other scenarios, our observations and those previously reported might reflect a deleterious effect of tau species present in the sarkosyl-insoluble fraction in cells close to the injection site as also suggested by [61]. We were able to determine this *in vitro* showing a relevant decrease of TUJ1 immunostaining after treatment with sarkosyl-insoluble fractions of P301S^+/−^ or AD, and *in vivo*. The negative effect of amyloid fibrils, prefibrils *vs* oligomeric forms in cell survival has been described for several amyloids including Aβ [15], insulin fibers [6], lysozyme fibrils [66] see also ([24] for additional examples and review) using several protocols and experimental approaches. In fact, although a clear consensus is still elusive, brain-derived sarkosyl-insoluble fractions might contain different forms of tau fibrils [80] that can interact with cellular membranes leading to membrane instability and increased toxicity (see [24] for review). If this is the case, these effects are PrP^C^ independent since the absence of its extracellular presence in neuronal parenchyma does not modify ptau presence and distribution in sarkosyl-inoculated mice. This scenario has been also described for other amyloid proteins that also render neurotoxicity in absence of PrP^C^ [53, 56, 116]. In conclusion, we consider that sarkosyl-insoluble fractions are not the appropriate tool to characterize the role of putative neuronal or glial receptors as PrP^C^ in the seeding and propagation of ptau. On the other hand, we tried to take profit from the extracellular presence of PrP^C^ to block seeding and spreading of tau. As indicated, the effects of the extracellular GPI anchorless PrP^C^ have been linked to a reduction of the toxicity of Aβ-oligomers [58, 69] and fibrillation [95], but do not affect folded Aβ [134]. Unfortunately, in our study its presence does not impair the seeding and phosphorylation of endogenous ptau after sarkosyl-insoluble AD fraction, most probably due to the large fibril content of the fraction, allowing the different pre-fibrillar species of tau to take effect on the recipient neurons.

In our study, PrP^C^ absence decreases the number of AT8-positive neurons in the CA1 of P301S transgenic mice and to a lesser extent in AAV2r.*Syn*.P301L hippocampal infection. A similar decrease in ptau was recently obtained by Stoner *et al.*, [119] although ptau analysis was developed in the retrosplenial cortex instead of the CA1 hippocampus in double mutant (*App*^NL−G−F^/*hMapt*) mice. In our case, we used the ZH3 *Prnp^0/0^* mice [97] with C57BL/6J background, and in Stoner *et al., Prnp^0/0^* mice were the Edinburgh strain [36, 88] with 129/Ola background. The expression of the mutated form of htau in the P301S mice is under the cellular prion protein promoter in a B6C3H/F1 background in contrast to C57BL/6J background of the (*App*^NL−G−F^/*hMapt*) mouse. Relevantly our data demonstrate a decrease of ptau in the absence of Aβ deposits, similarly, as happens in the above-cited manuscript with Aβ presence. Although differences in genes affected by the absence of PrP^C^ have been described in several manuscripts with differences in the genetic background (i.e, [14, 28, 103], we previously reported that ZH3 *Prnp^0/0^* mice showed decreased levels of tau and increased 3R/4Rtau ratio *vs* wild-type mice [82]. In addition, changes in ptau and levels of tau were described during the development of FVB/N *Prnp^0/0^* mice *vs* wild-type [14]. Although a clear explanation of whether the absence of *Prnp* might decrease ptau is lacking in both studies, our present data reinforce the notion of an active role of PrP^C^ in decreasing ptau generation independently of Aβ, emphasizing the data that PrP^C^ could be a putative target for future therapeutic strategies in tauopathies.

## Data Availability Statement

The raw data supporting the conclusions of this article will be made available by the authors upon request, without undue reservation.

## Author Contributions

JS-J, VG, PA-B, and LL performed and designed experiments and performed data analysis. FH, JA, MN, AA, IF, EYH, JS and CFLL and RG designed experiments and edited the manuscript. JADR and JS provided experimental funds and edited the manuscript. All authors contributed to the article and approved the submitted version.

## Acknowledgements

The authors thank Tom Yohannan for his editorial advice. We also thank Miriam Segura-Feliu and Juan José López for their technical help. We would also like to thank Oscar Castaño for the help in developing *ad hoc* Python 3.9 based graphic user interphase (GUI) for calcium analysis and our collaborators in providing Matlab^TM^ algorithms and reagents for this study. We also thank the microFab nanotechnology platform of IBEC for their technical help. The authors also thank all the members of the Del Río, Ferrer, Ávila, and Aguzzi laboratories for their helpful comments.

## Supplementary information

**Suppl. Figure 1.**
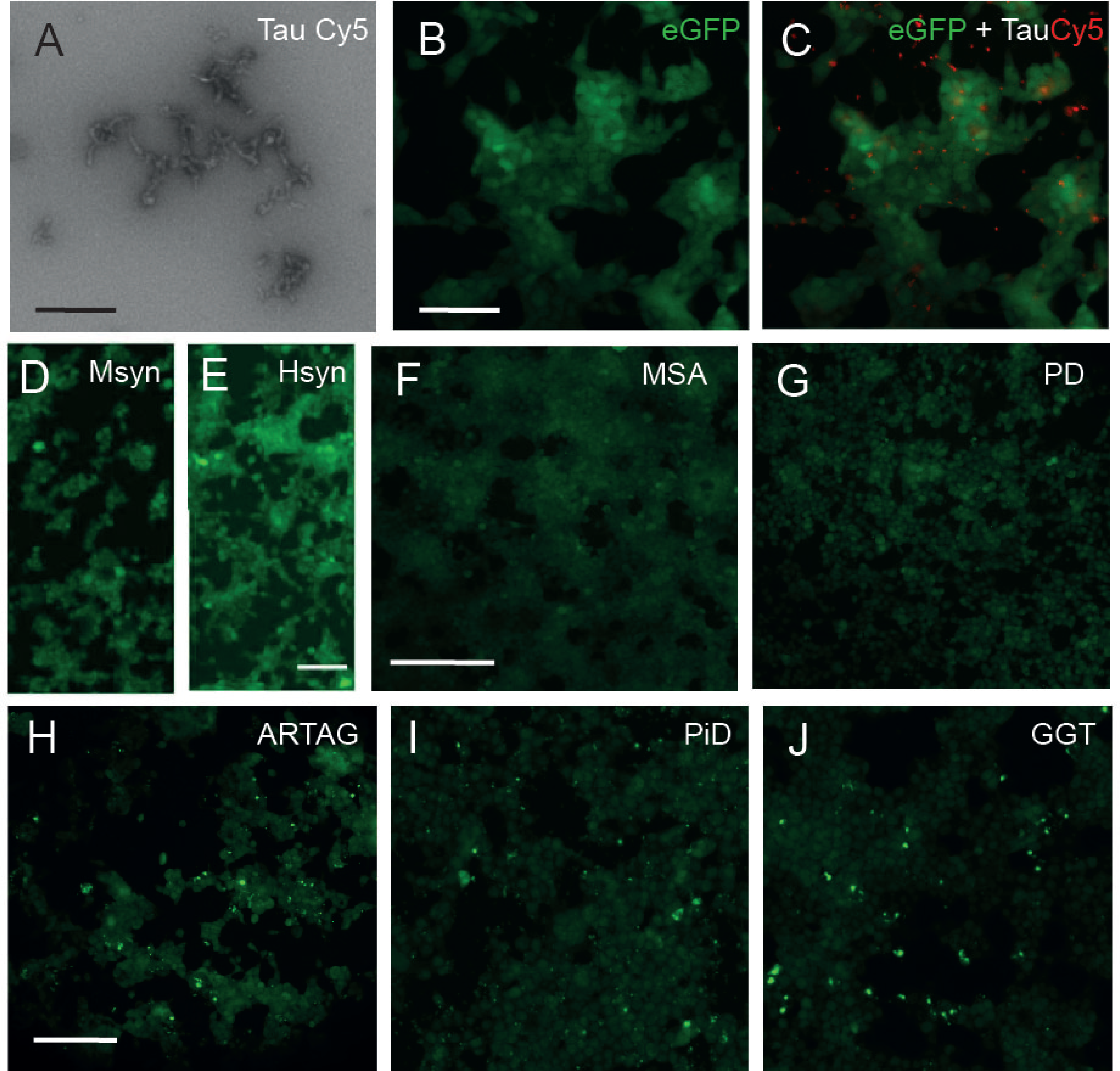
Seed specific fluorescence revealed in the Tau RD P301S Biosensor cell line. **A)** TEM image of the tau-Cy5 used in the experiment. **B-C)** Tau Cy5 can be uptaken by the Biosensor cell line but is unable to induce aggregation in the reported cells. **D-G)** Absence of fluorescence aggregates in the Biosensor cell line after the incubation with m-syn (D), h-syn (E) prefibrils, sarkosyl extracts from MSA (F) and PD (G). **H-J)** Formation of fluorescence aggregates after the incubation of the Biosensor cell line with sarkosyl-insoluble extracts from ARTAG (H), PiD (I) and GGT (J). Scale bars: A = 1 μm; B = 100 μm pertains to C; D-E = 50 μm; F = 200 μm pertains to G; H = 200 μm pertains to I-J.

**Suppl. Figure 2.**
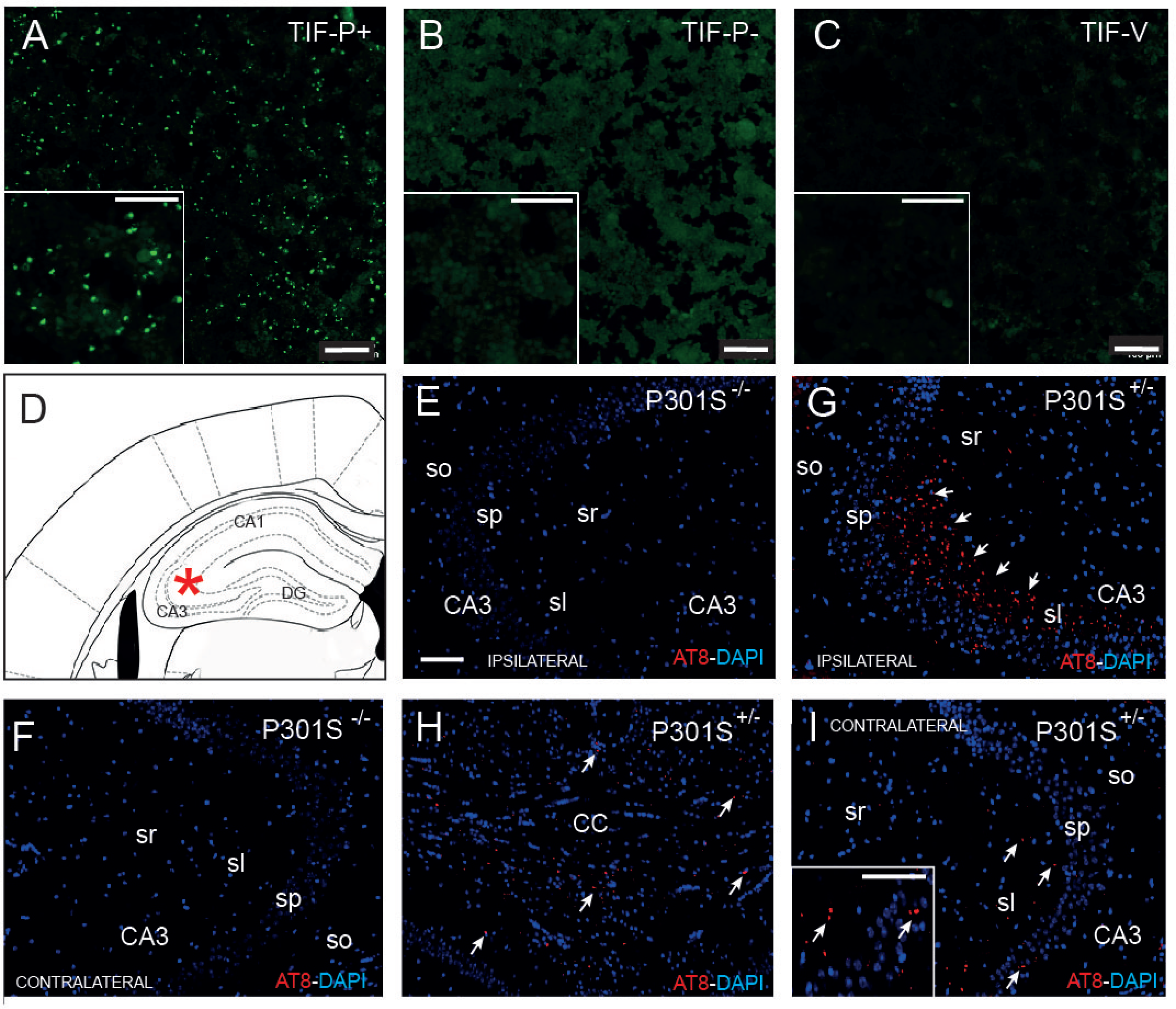
Seed properties of the aggregates generated in the Biosensor cell line. **A-C)** Freshly prepared Biosensor cell cultures were incubated with X-Triton extracted aggregates (TIF-P+, TIF-P-, and TIF-V; see Results for details). Only TIF-P+ was able to induce *de novo* formation in this second passage experiment. **D)** Scheme of the location of the inoculation of the X-Triton extracted TIF-P+ aggregates. **E-I)** Results of the inoculation of TIF-P+ in non-transgenic (E-F) and transgenic (G-I) P301S mice. TIF-P+ aggregates were able to induce AT8-positive ptau deposits in P301S transgenic mice (ipsilateral (G) in corpus callosum (H) and contralateral (I) hippocampus) but not in non-transgenic mice (E-F). The insert in I is a higher magnification of AT8-positive deposits in the contralateral hippocampus. Abbreviations as in Figure 2. Scale bars: A-C = 50 μm; E = 100 μm pertains to G-I; Insert in J = 75 μm.

**Suppl. Figure 3.**
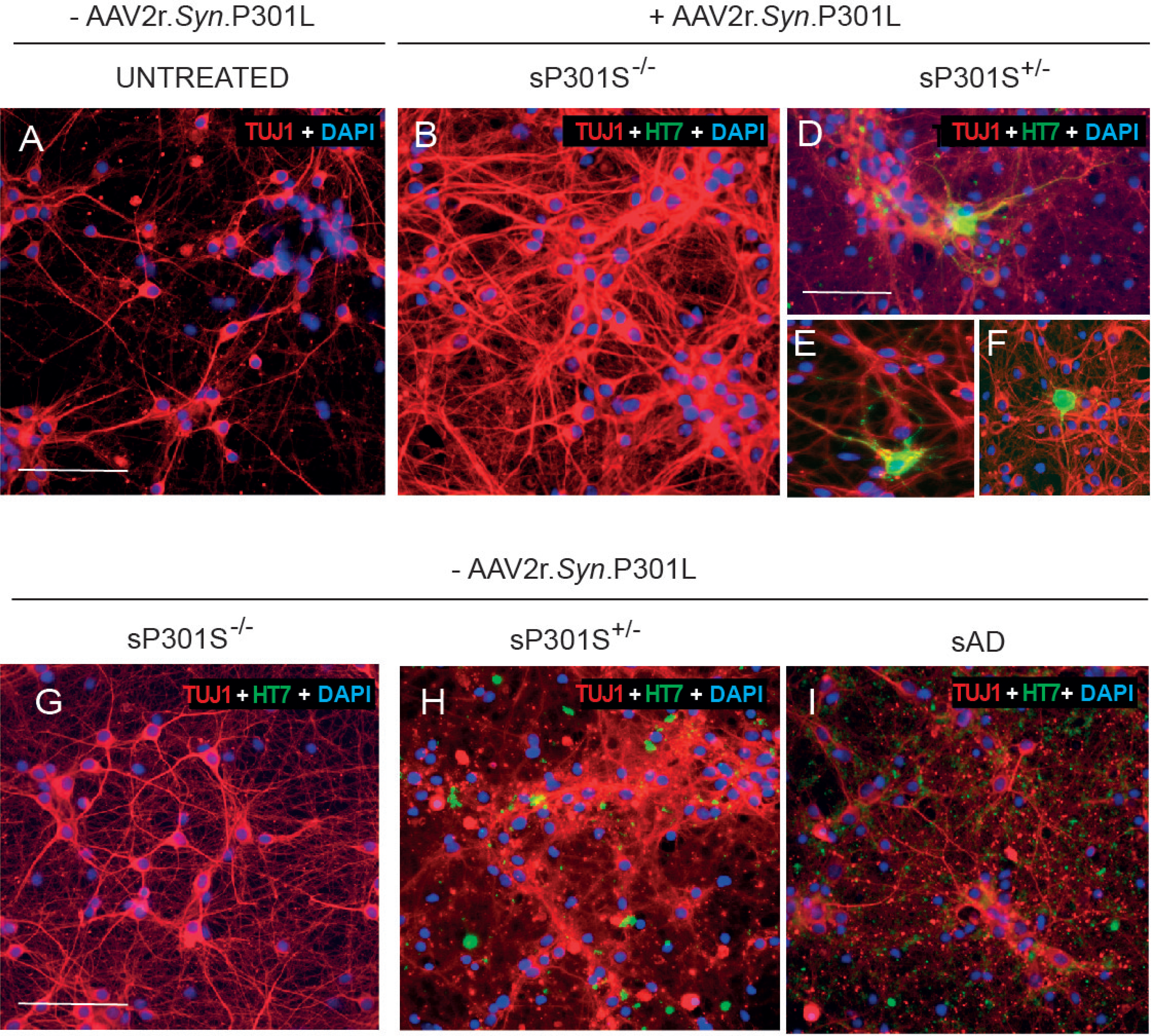
Effect of the endogenous expression of P301L in human tau aggregation after sarkosyl-insoluble treatment is primary cortical neurons. **A-F)** Primary cortical neurons from wild-type mice were uninfected (A) or infected with AVV2r.*Syn*.P301L viruses (see Material an Methods for details). Afterwards, cultures were double processed for HT7 and TUJ1 immunodetection. In contrast to untreated or incubated with sP301S^−/−^, cells treated with sP301S^+/−^ sarkosyl fraction showed double labeled neurons (D-F). In a second set of experiments untreated cells were incubated with sP301S^−/−^ (G); sP301S^+/−^ (H) or sAD (I). After incubation cultures were fixed with Methanol and immunostained as in (B-F). Relevantly, the immunostaining against TUJ1 renders dystrophic and fragmented neurites indicating ongoing cytotoxicity. Scale bars: A = 75 μm pertains to B; D = 50 μm pertains to E-F; G = 75 μm pertains to H-I. Immunoreacted cultures were counterstained with DAPI.

**Suppl. Figure 4. AD-derived sarkosyl-insoluble inoculated mice showed AT8-P62 double labelled cells.** A-E) Fluorescence photomicrographs illustrating examples of *Prnp^+/+^* (A-C) and GPI^−^ *Prnp* (D-E) immunoreacted sections using antibodies against P62 and AT8 at 3 post inoculation. Numerous double labelled cells can be observed in both phenotypes (arrows in A-E). Scale bars: A = 100 μm pertains to B-E.

**Suppl. Table 1.** Statistical analysis of the histograms of Figure 7F-G

**Suppl. Videos 1-3.** Examples of a control primary culture infected with GCaMP6s and analyzed at day T0 (Suppl. Video 1); T3 (Suppl. Video 2) and T7 (Suppl. Video 3). Video frame rate: 25 fps, one frame each 500 ms.

**Suppl. Videos 4-6.** Examples of a ptau treated primary culture infected with GCaMP6s and analyzed at day T0 (Suppl. Video 4); T3 (Suppl. Video 5) and T7 (Suppl. Video 6). Video frame rate: 25 fps, one frame each 500 ms.

**Suppl. Videos 7-9.** Examples of a tau treated primary culture infected with GCaMP6s and analyzed at day T0 (Suppl. Video 7); T3 (Suppl. Video 8) and T7 (Suppl. Video 9). Video frame rate: 25 fps, one frame each 500 ms.

**Suppl. Videos 10-11.** Examples of a ptau treated primary culture (200 nM) infected with GCaMP6s and analyzed at day T0 (Suppl. Video 10); and T3 (Suppl. Video 11). Video frame rate: 25 fps, one frame each 500 ms.

**Suppl. Videos 12-13.** Examples of a tau treated primary culture (200 nM) infected with GCaMP6s and analyzed at day T0 (Suppl. Video 12); and T3 (Suppl. Video 13). Video frame rate: 25 fps, one frame each 500 ms.

